# Unveiling the Ecology of *Legionella* in a Stratified Freshwater Lake: Seasonal Dynamics, Host Interactions, and Persistence Under Oxygen Limited Conditions

**DOI:** 10.1101/2024.12.02.626374

**Authors:** O. Berman, Y. Be’eri-Shlevin, S. Ninio

## Abstract

**Abstract:** *Background:* *Legionella* are predominantly recognized as aerobic pathogens in man-made water systems. However, their potential persistence in natural freshwater environments, particularly under oxygen limited conditions, remains poorly explored. In this study we investigated the spatio-temporal dynamics of *Legionella* occurrence in a seasonally stratified sub-tropical freshwater lake, with a focus on anaerobic conditions of the anoxic hypolimnion.

*Results:* Our study reveals significant seasonal variations in *Legionella* absolute abundance, with the highest concentrations occurring during and immediately following lake mixing events. Unexpectedly, high levels of *Legionella* were measured in the anaerobic hypolimnion layer of the lake. Utilizing genus specific amplicon-based sequencing, we found significant shifts in *Legionella* community composition, that are related to the sampling month. Several environmental factors were associated with the observed changes, including temperature, DO, chlorophyll and dinoflagellate biomass. Moreover, we identified *Legionella* genotypes unique to samples presenting hypoxic conditions - that were not closely related to known *Legionella* species. In addition, we noted genotypes present in anoxic samples, that were absent from the oxic layers of the corresponding sampling dates. These results were accompanied by changes in the interaction patterns between *Legionella* and their potential hosts, in oxic and anoxic conditions.

*Conclusions:* This study challenges the conventional view of *Legionella* as a strictly aerobic pathogen by demonstrating its persistence in anoxic freshwater environments. Our findings suggest that certain *Legionella* species may have adapted to low- oxygen conditions, potentially through alternate metabolic pathways or by residing within protozoan hosts. The identification of unique *Legionella* genotypes in the hypolimnion, along with shifts in occurrence, diversity, and host interactions, underscores the complexity of *Legionella* ecology. These results highlight the need for further research on *Legionella* in natural freshwater systems, which serve as reservoirs for the bacteria and potential sources for human infection. Further investigation into the mechanisms underlying *Legionella* persistence in anaerobic conditions and its interactions with environmental hosts is essential for a better understanding of the evolutionary forces shaping this family of human pathogens.

## Introduction

*Legionella* is a gram-negative bacterium [1], initially isolated in Philadelphia following an outbreak in the 1976 American Legion Convention [2,3]. *Legionella* infections are responsible for the majority of USA waterborne related outbreaks and hospitalizations, with the latter estimated at between 8,000-18,000 cases annually [4,5]. The majority of outbreaks are sporadic and considered to be non-related [6]. Members of the *Legionella* genus are ubiquitously found in a variety of aqueous environments, both natural and man-made. These include lakes, rivers, ponds, hot springs, wet soil, hot water systems, cooling towers and sewage [7–13]. Concomitantly, *Legionella* can persist and grow in a wide range of physical and chemical environmental conditions (e.g., temperature, dissolved oxygen (DO), pH, salinity, iron concentrations and dissolved organic matter) [8,9,11,14,15]. Traditionally, *Legionella* has been considered an obligate aerobe, with studies focusing on *Legionella* occurrence in natural water bodies and controlled laboratory experiments [7,8,16–20]. This view is widely accepted to date, despite the fact that the number of studies assessing the effect of dissolved oxygen (DO) concentrations on *Legionella* growth, persistence and natural occurrence is limited [21]. Most of our knowledge regarding the environmental conditions required for the growth of *Legionella* spp. comes from lab experiments using clinical isolates, many of them are *L. pneumophila* serogroups. In these settings, the *Legionella* genus is considered fastidious, with isolation and growth possible only on highly permissive media, containing among others, L-cysteine, alpha-ketoglutarate, ferric pyrophosphate and yeast extract [22,23]. Paradoxically, in natural environments, *Legionella* are frequently detected in oligotrophic conditions [14,24]. This discrepancy in growth requirements, coupled to the low nutrient content present in oligotrophic water, is explained by their ability to persist in biofilms [25,26] and by reported associations of these bacteria with protozoa. Intracellular replication has been demonstrated in-vitro within amoeba, ciliates and slime mold [27–33] and other associations reported with a number of cyanobacteria species [14,34–36]. For example, when *L. pneumophila* and *Fischerella sp.* are co-cultured, the former apparently use photosynthetic substances and possibly other extracellular products, produced by the cyanobacteria, as carbon and energy sources [37]. It has been demonstrated that *Legionella* employs the same genes to replicate within both *Acanthamoeba castellanii* and human macrophages [38].

The dynamics of *Legionella* spp. diversity and community composition have been investigated mainly for *L. pneumophila* in man-made structures, typically reporting higher levels during summer. These include, among others, water distribution systems, cooling towers, potable water systems and grey water systems [39–41]. Epidemiological studies performed in various countries (e.g., Scotland, Korea, Japan, USA and Israel) point to a more complex and variable seasonal pattern [42–46]. Yet overall, the data points to elevated rates of infection during summer. The latter has been associated with higher precipitation, humidity, temperature and heavy rainfall [47–49]. However, the data heretofore depict human infection rates and thus relevant mainly to *L. pneumophila* in man-made facilities and does not necessarily reflect *Legionella* spp. in natural environments. Studies on the latter are relatively scarce and until recently have been restricted to cultivation and infection methods [8,50–52], limiting the ability to represent the seasonal abundance of the genus as a whole. Indeed, the development of cultivation-independent techniques enabled us to inspect the natural occurrence, abundance, diversity, community composition and environmental conditions related to *Legionella* spp. [53–59]. However, despite their frequent detection in natural aquatic habitats and potential function as a natural pathogenic reservoir that may leak into man-made facilities, only a small number of studies focused on the natural occurrence of *Legionella* spp.

As a result, *Legionella* diversity in natural environments is underestimated, as is evident by the identification of unclassified *Legionella* distinct from known representatives in culture independent techniques [60]. In addition, better understanding of the environmental conditions influencing abundance and seasonality of *Legionella*, can aid in their control in man-made surroundings.

The main aim of this study is to better understand the ecology of *Legionella* spp. in a natural freshwater lake environment. We sought to delineate the occurrence of *Legionella* spp. and their potential hosts, and to understand which environmental conditions affect spatio-temporal variations in their abundance, diversity, and community dynamics under oxic and anoxic conditions. To this end, we followed the monthly occurrence and community composition of *Legionella* spp. in a sub-tropical lake, alongside both biotic and abiotic parameters over the course of two and a half years, providing the first comprehensive analysis of *Legionella* dynamics in a stratified lake ecosystem. Our results challenge the long-held view of *Legionella* as an obligate-aerobes and fill critical gaps in our understanding of the ecology and evolution of this family of pathogens in the natural environment.

## Materials and methods

### Study site

We studied Lake Kinneret (Sea of Galilee) a sub-tropical monomictic, mesotrophic freshwater lake, located in the northern part of the of the Afro- Syrian-Rift, northern Israel (Fig. 1). The lake Spans approximately 170 km2, with an average depth of 24m and a maximal depth of 43m. During April-May to December, the lake is stratified and divided into thermally and chemically distinct layers (i.e., the epilimnion, metalimnion and hypolimnion). The first two are warmer (24–30 °C) and oxygenized, while the latter is anoxic and cooler (14–16 °C). Situated at the center of the lake, (32°82.146′N - 35°35.191′E, ∼40m depth) station A is located in the vicinity of its deepest point and is routinely monitored year long.

**Figure 1.**
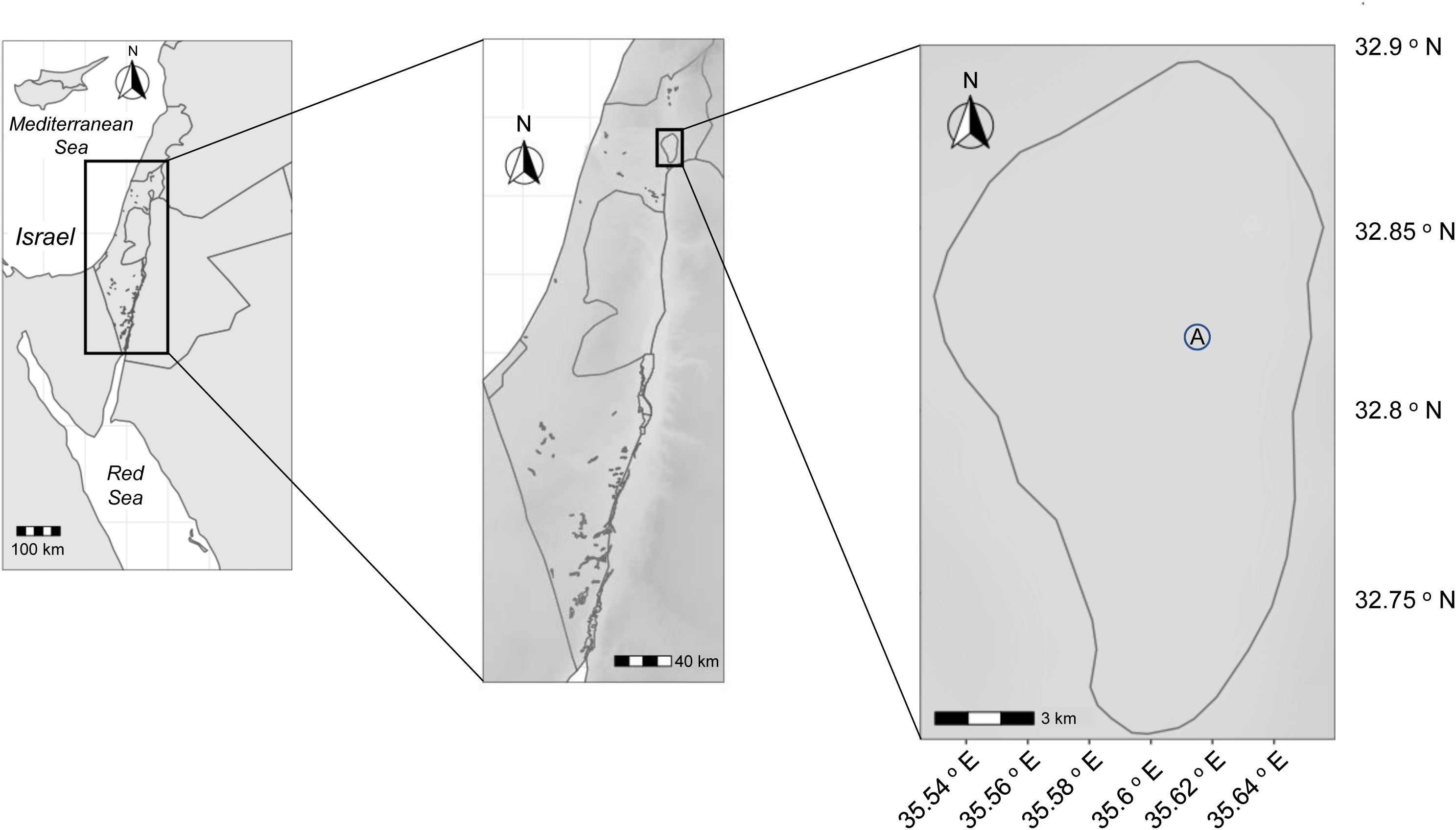
Map of study site. All samples were collected at station A (32°82.146′N - 35°35.191′E, ∼40m depth), located at the center part of the sea of Galilee (right panel). This warm monomictic, mesotrophic freshwater lake, is located in the northern part of the of the Afro-Syrian-Rift, northern Israel (middle panel), the Middle East (left panel).

### Sample collection, preparation and physicochemical analysis

Monthly samples were taken at station A, between May-November 2017 at two depths (1m and 25m), and between December 2017 to December 2019 at three depths - 1m, 15m and 25m (n=84). We determined the identity of sampling layer based on sampling depth and the vertical location of the thermocline (Fig. 2A lower panel). In addition, extended depth profiles were taken during March 2018 (1, 3, 5, 10, 15 and 25m) and June 2018 (0, 1, 2, 3, 5, 7, 10, 13, 14, 15, 16, 17, 18, 19, 20, 22 and 25m). Samples were taken using a vertical Niskin Water Sampler (duplicates, stored in 1-L sterile plastic bottles) and transferred to the laboratory within 2-3 hours. Approximately 500 μl were filtered on 47 mm diameter 0.22 μm pore size, Supor Polycarbonate Membrane Disc Filters, (Pall, MI, USA) and transferred immediately to -80°C until further use. 1 L of water were filtered through a 0.45-mm membrane filter (Sartorius, Göttingen, Germany) and used for chemical analysis.

**Figure 2.**
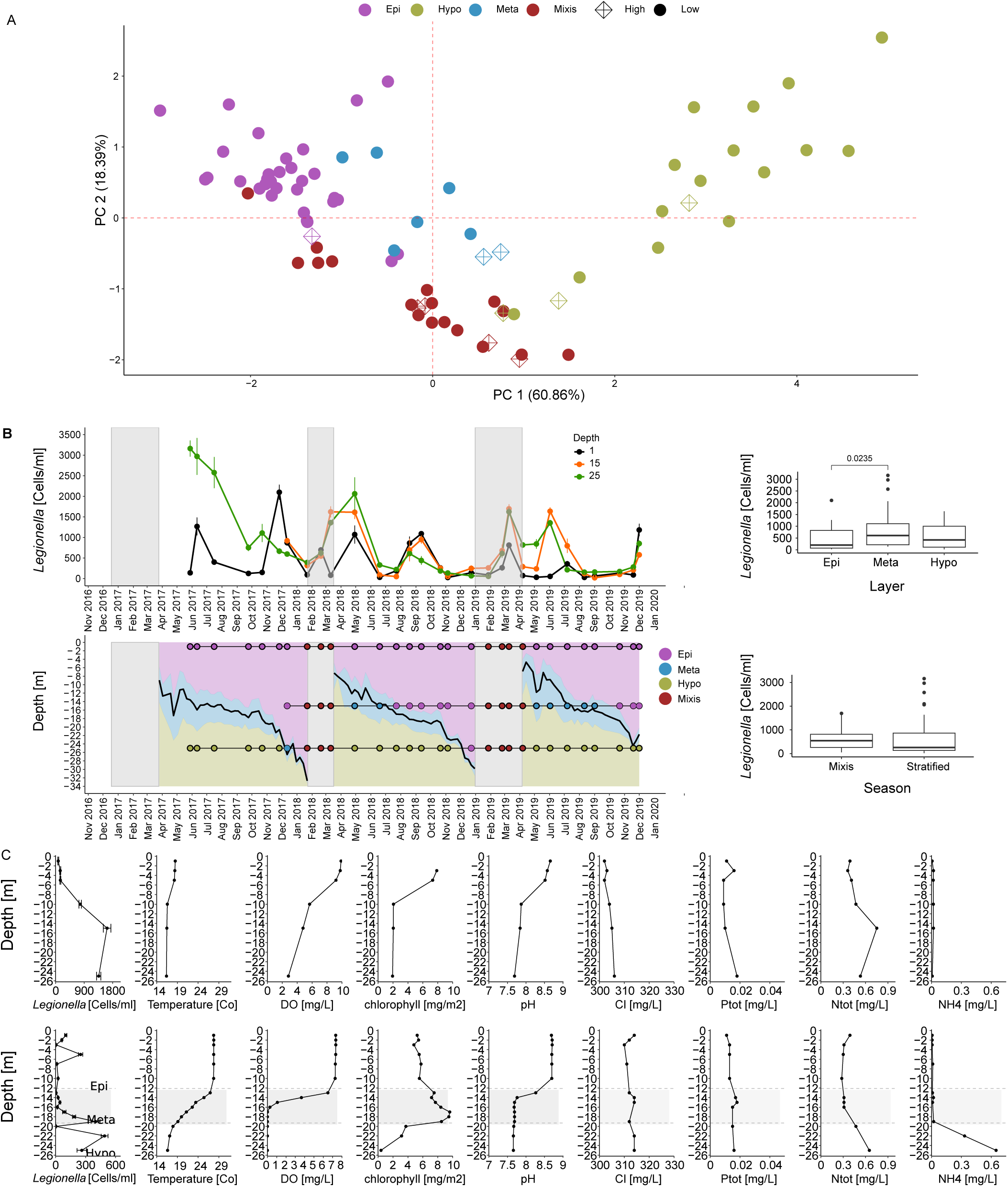
Station A chemical characteristics and *Legionella sp.* absolute abundance levels. **(A)** Principal component analysis (PCA, n=86) was performed for selected physicochemical parameters measured at station A, from May-2017 to December-2019. The PCA included the following environmental variables: Temperature, DO, SO4^2-^, pH, NH4^+^ and chlorophyll. Loading scores and vectors are presented in Table S11 and Figs. S2. The environmental variables were chosen following a PCA analysis from a larger subset of environmental factor (see Materials and Methods, Fig. S1A-S1B and Tables S9-S10). Each sample is denoted by a circle, and colored according to the layer sampled, as indicated. Samples with high values of *Legionella* absolute abundance are marked with a diamond (the cutoff was determined as values above the average + SD). **(B)** *Legionella* spp. specific primers and probes were used to quantify absolute abundance levels (upper left panel) at station A at 1m, 15m and 25m. Sampling dates (May 2017 to December 2019) and position in the water column are presented (bottom left panel). Samples were assigned to the relevant color, indicating the distinct water layer. During stratification: Epilimnion (Epi=purple, n=34), metalimnion (meta=pale-blue, n=8) and hypolimnion (Hypo=mustered-green, n=21), or to complete water column mixing (mixis=maroon, n=21). Samples were denoted by point indicating the relevant layer, at the appropriate sampling depth (1m, 15m and 25m; denoted by black lines at the relevant depth). ANOVA and post-hoc tukey tests were performed to analyze differences between sampling layers (upper right panel, n=63, Table S1), and a t-test was performed to access the differences between the stratified and mixis preriods (bottom right panel, n=84, Table S4). **(C)** Extended depth profiles taken during the mixis (March 2018, upper panel) and stratification (June 2018, lower panel) periods. For both dates, depth profiles of *Legionella* spp. absolute abundance levels, temperature, Dissolved Oxygen (DO), chlorophyll, pH, Cl, total phosphate (Ptot), total nitrogen (Ntot), and NH4 are presented.

### Environmental parameters

The physicochemical parameters: Temperature, Dissolved Oxygen (DO), pH, conductivity, turbidity and Chl were measured in situ using an autonomous profiling unit with a Manta+ multiparameter probe (Eureka, Austin, Texas, USA), sampling at intervals of 0.5m and 6hr. The chemical parameters alkalinity, Cl, total nitrogen (Ntot), total phosphorus (Ptot), Cl, Ca, SO4^2+^, NO3^−^ and NH4^+^, were analyzed following standard methods (APHA 2005) and are further described by Nishri Nishri (Nishri, 2011).

### DNA extraction

DNA extraction was performed on approximately 300 μl water, using the power water DNA isolation kit (QIAGEN, Hilden, Germany; prior MoBio, CA, USA), according to the manufacturer’s instructions with the following exceptions: Alternative lysis method was performed. After the addition of lysis buffer, samples were incubated at 65°C for 10 min. Elution buffer was pre- warmed at 70 °C for 5 min, to increase DNA recovery. Eluted DNA was precipitated with sodium acetate. Briefly, 0.1 volume sodium Acetate (3 M, pH 5.2) was added to each sample and vortexed. X3 volumes of 100% pure ethanol (Merck) was added and incubated at room temperature for 15 min. Following centrifugation (14,000g, 30 min, 4°C), the pellet was rinsed in 70% pure ethanol, centrifuged again and transferred to -80°C until use.

### Real time PCR (qPCR)

Quantification of *Legionella* spp. was performed on DNA extracted from all station A water samples (see sample collection and preparation), as previously described[62]. Briefly, qPCR for the 16S gene (n=84) was done, using primers Leg- F1c (TAGTGGAATTTCCGGTGTA), Leg- R1c (CCAACAGCTAGTTGACATC) and the probe ([6FAM]-CGGCTACCTGGCCTAATACTGA-[BHQ1]). The reaction was performed using the TaqMan™ Fast Advanced Master Mix (Applied Biosystems, CA, USA). The reaction mix (20 μl) contained: 2 μl DNA template, 10 μl of x2 qPCR buffer, 3 μl each primer, 1 μl TaqMan probe (both by Sigma-Aldrich, Israel; to final concentrations of 450 nm and 250 nM, respectively) and 1 μl molecular water (Biological Industries, Israel). Reaction conditions included 2 min of incubation at 50 °C followed by 10 min of activation at 95 °C and 40 cycles of denaturation 15 sec at 95 °C and a combined annealing/extension step of 60 sec at 50 °C. Gene concentration was determined based on calibration curves of standards with known concentration of the target gene vs cycle threshold (CT), using Rotor Gene 6000 software version 1.7 (Corbett Research, UK).

### Biomass counts

For a random forest analysis (described later in the method section in detail), we evaluated total ciliates and phytoplankton biomass of specific lineages (Fig. S3B) were evaluated. In addition, the biomass of major phytoplankton taxonomic groups was measured: Dinophyta (Dinoflagellates), Chlorophyta (green algae), Cryptophyta (cryptophytes), Bacillariophyta (diatoms), Cyanophyta (cyanobacteria) and *Microcystis sp.*, as previously described [63,64]. Briefly, water samples from station A (at 1m) were Lugol-preserved and microscopically counted by the sedimentation chamber technique (Lund et al. 1958). Nanoplanktonic (10 ml) and large species (1 ml) were sedimented for 24h. Biomass calculation was performed by multiplying counts per ml with average biovolume, assuming a specific density of one [65]. Both parameters were evaluated by a specialized computer program, PlanktoMetrix.

### Bacterial productivity

Bacterial productivity was estimated using the 14C-leucine incorporation method, essentially as described before [66]. Briefly, Samples (1.7 ml) were incubated with 235 nM leucine (Revvity, specific activity 328 Ci mmol–1) for 3h under ambient temperature. Triplicate additions of trichloroacetic acid (TCA) were performed for each sample, before 14C-leucine introduction, as controls. The incubations were terminated with 92 μL of concentrated cold TCA (100%), followed by micro centrifugation and 5% TCA wash of the precipitated cells. After the addition of 1 mL scintillation cocktail (Ultima-Gold) to each vial, the samples were counted using a TRI-CARB 2100 TR (Packard) liquid scintillation counter.

### PCA analysis

We sought to characterize the differences in various environmental conditions related to both sampling layer and sampling season (mixis vs. stratified periods). We performed an initial PCR analysis (n=86) on the water samples obtained between May 2017 to December 2019 (Figs. S1A-S1B and Tables S9-S10). We initially included 14 environmental parameters, and omitted the ones showing strong cross-correlation. The initial PCA analysis included 10 environmental parameters: Temperature, DO, Cl, SO4^2-^, Turbidity, pH, NH4^+^, Nitrite, Ptot and Chl (Table S9 and Supplementary data file 2). After omission of variables presenting weak Loading scores (Table S10), a final PCA was performed with 6 environmental parameters: Temperature, DO, SO4^2-^, pH, NH4^+^ and chlorophyll. In addition, the data was separated by stratified and mixis periods. Prior to PCA analysis all samples were centered and scaled (log+1 values).

### Random forest

To determine which environmental parameters were related to *Legionella* abundance, we performed a machine learning procedure (random forest analysis). *Legionella* absolute abundances were regress against the aforementioned abiotic (n=84) and biotic (n=30) parameters in two separate analyses, using packages caret [67] and randomForest [68]. For the biotic analysis only samples from 1m (epilimnion) were included, as biotic data was only available for this depth. The random forest models (ntrees=1001) were run with the mtry parameter set to default and error was measured as out-of-bag (OOB) error. Root Mean Squared Error (RMSE) was used to assess prediction accuracy.

### Next Generation Sequencing (NGS)

To explore the microbial community of *Legionella* spp., we utilized NGS amplicon-sequencing, performed on water samples (n=28) collected during: September 2017, January, March, April, June and September 2018 and January, March, June and December 2019. September 2017 and January 2018-2019 samples were of 1m. All other samples of 1m, 15m and 25m. 16S rRNA gene amplicon-sequencing of the V3-V4 variable regions, was done using specific *Legionella* spp. primers: 5′- GGCCTACCAAGGCGACGATCG-3′ (Lgsp17F, forward) and 5′- CACCGGAAATTCCACTACCCTCTC-3′ (Lgsp28R, reverse), as previously described [69], sequenced on an Illumina MiSeq V3 platform, yielding 2×300 bp paired-end reads. We have previously shown that use of this genus specific primer set is superior to universal primers for the identification of *Legionella* representatives [70]. To examine potential *Legionella* hosts, NGS amplicon-sequencing designed for protozoa [71] was performed on the 18S rRNA gene targeting the V9 region utilizing primers: 5’- GTACACACCGCCCGTC-3’ (EUK1391F, forward) and 5’-CCTTCYGCAGGTTCACCTAC-3’ (EUK1510R, reverse), on an Illumina MiniSeq platform, yielding 2 × 153 paired-end reads following the manufacturer’s guidelines. Library preparation and sequencing of the September-2017 16S DNA extracts were performed at the MR DNA sequencing center (www.mrdnalab.com, Shallowater, TX, USA). Sequencing was done with barcodes on the forward primer, in a 30-cycle PCR reaction using the HotStarTaq Plus Master Mix Kit (Qiagen, USA). PCR conditions for the *Legionella* specific primers were 94 °C for 3 min, followed by 30 cycles of 94 °C for 30 sec, 68 °C for 40 sec and 72 °C for 1 min and a final elongation step at 72 °C for 5 min. For the 18S universal primers, the same conditions were used, with an annealing temperature of 57 °C. PCR products were visualized on a 2% agarose gel to determine bands relative intensity. Pooled samples were purified using calibrated Ampure XP beads. Then the pooled and purified PCR products were used to prepare an Illumina DNA library. Raw sequence data were analyzed as previously described [72]. First, MacQIIME 1.9.1 [73] was used to achieve proper read orientation of sequence reads obtained from MR DNA sequencing center, using qiime-extract.barcodes.py. The raw sequences were sorted based on orientation and barcodes were extracted. Raw sequence data were then processed using QIIME2 2020.11 [74]. Data was demultiplexed and quality filtered using the q2-demux plugin. Denoising was performed with DADA2 [75], using the q2-dada2 plugin. We examined Amplicon Sequence Variants (ASV) and not Operational Taxonomic Units (OTUs), as it enables distinguishing single nucleotide variations [76]. To account for length variations, Amplicon Sequence Variants (ASVs) were defined by clustering at 100% similarity [77]. For each primer set, all independent runs were merged post-denoising, ASVs occurring under a total frequency of 20 reads in less than 2 samples were removed. Taxonomy analysis was done with the 138-SILVA database QIIME release (at 99% clustering) [78], and the classifier was generated using the q2-feature-classifier plugin [79], using the extract-reads and fit-classifier-naive-bayes methods. The classifier generated was then used for the classification of the ASVs using the classify-sklearn method (ver. 0.23.1) [80]. To study the microbial populations of *Legionella* spp., we filtered the feature-frequency tables to retain 16S ASVs classified to *Legionella* spp. only (Supplementary Data 3). As SILVA classification is largely successful up to the genus level, *Legionella* ASVs were left unclassified for most analyses. Phylogenetic analysis was performed with reference *Legionella* sequences (see Phylogenetic analysis section below). Potential *Legionella* spp. hosts were assessed by filtering the 18S ASVs to retain only those classified as Amoebozoa, Ciliophora (Ciliates), Percolozoa (Excavata) and Dinoflagellates (Supplementary Data 5). All analyses were performed on these filtered tables. Alpha (Shannon’s entropy, Faith’s PD and Pielou’s evenness) and beta (Jaccard similarity coefficient and Bray-Curtis dissimilarity indices) diversity analyses were assessed at rarefying depths of 406 and 6424 reads per-sample for the 16S and 18S rRNA genes, respectively (using the core-metrics-phylogenetic function of the q2-diversity plugin [81]. Beta-diversity was visualized using the q2-emperor plugin. To study *Legionella* diversity according to sampling month, while maintaining a relatively balanced study design, we included in the analysis only the months that were sampled at all three depths (n=21). Rarefaction curves were generated at varying depths to ensure ASVs identification reached a plateau and to justify the rarifying depth chosen for the diversity analysis (using the alpha-rarefaction function of the QIIME2-diversity plugin, Figs. S4 and S9). Statistical analysis for all indices was done using the alpha-group-significance (Kruskal-Wallis tests, p-values adjusted according to the Benjamini-Hochberg FDR correction) and beta-group-significance (PERMANOVA, PERMDISP and ADONIS with 999 permutations, p-values adjusted according to the Benjamini-Hochberg FDR correction). Un-rarefied frequency filtered tables of absolute counts, and matching tables converted to relative abundance (using the relative-frequency function of the q2-feature table plugin), were exported in biome format and used for downstream statistical analysis and visualization in R, using the following packages: Phyloseq [82], ggplot2 [83], VennDiagram [84]. Heatmaps were created using the q2-feature-table heatmap plugin.

### Phylogenetic analysis

Phylogenetic trees were constructed for *Legionella* spp. ASVs obtained from the NGS amplicon-sequencing, together with previously characterized reference *Legionella* sequences denoted with the corresponding NCBI accession numbers (Supplementary Data 4). A phylogenetic tree was created for abundant ASVs (>15%). In addition, unfiltered trees (min 20 reads, in at list 2 sample) were constructed for month and sampling layer (n=28 samples). The reference sequences were manually trimmed to match the length of the ASVs. Evolutionary analyses were conducted in MEGA11 version 11.0.9 [85], alignment of all ASVs was carried out using the imbedded muscle algorithm [86]. The evolutionary history for all trees was inferred using the Maximum-Likelihood method and Hasegawa-Kishino-Yano (HKY) [87] model. *Coxiella burnetii* was included as an outgroup. Initial trees for the heuristic search were obtained automatically by applying Neighbor-Join and BioNJ algorithms to a matrix of pairwise distances estimated using the Maximum Composite Likelihood (MCL) approach, and then selecting the topology with superior log likelihood value. Bootstrap values (1000 replicates) above 50% are denoted by a black dot next to the branch. Trees are drawn to scale, with branch lengths indicating the number of substitutions per site. Trees were visualized using the R package ggtree [88].

### Cross-domain co-occurrence networks

To study the associations between *Legionella* ASVs and their potential protozoan hosts, we investigated the cross-domain co-occurrence patterns, based on the described above 16S and 18S rRNA NGS amplicon-based sequencing data. (Jiang et al., 2019; Matchado et al., 2021; Weiss et al., 2016) *Legionella* raw data were separated to samples with either oxic, or anoxic conditions (minimal total frequency of 20 reads, in a minimum of 2 samples) ASVs. We also filtered the 18S data to only contain known potential protozoan hosts (i.e., Ciliophora, Amoebozoa, Percolozoa and Dinoflagellates; a minimal total frequency of 20 reads, in a minimum of 2 samples). Correlation networks were constructed via the SPIEC-EASI[89] R package version 1.1.2, as described by Tipton et al 2018 [90] and https://github.com/zdk123/SpiecEasi. Normalization was performed internally within the spiec.easi function, by first converting the data to relative abundance followed by center-log scaling. We utilized the sparse graphical lasso (glasso), with optimal sparsity parameter based on the Stability Approach to Regularization Selection (StARS), threshold set to 0.1. Networks were analyzed using functions of the R package igraph[91] version 1.2.6. Edges with low weights (>0.01) and unconnected nodes were removed and only positive associations between ASV nodes are presented. Edge width corresponds to associations strength between ASVs.

### Statistical analysis and LMM model fitting

Statistical analyses were performed using the R (v. 4.0.3) rstatix package and the QIIME2 (v. 2020.11) software. Linear mixed effect model (LMM; lme4 package in R) was used to assess the *Legionella* absolute abundance variations between sampling months (fixed effect), while controlling for variations related to layer (random effect). Variables were log transformed, scaled and centered to meet the model assumptions. Normality was assessed by the Shapiro–Wilk test (p-value > 0.05). Formula: Logscale ∼ sampling month + (1 | layer). Comparisons of *Legionella* spp. absolute abundance between the different sampling layers were analyzed by the ANOVA and post-hoc tukey tests. Differences between stratified and mixis period were performed by t-test. Linear regressions were performed by the spearman method. Statistical analysis for all microbiome diversity indices was done using the alpha-group- significance (Kruskal–Wallis tests) and beta-group-significance (PERMANOVA, PERMDISP, and ADONIS with 999 permutations). p-values were adjusted according to the Benjamini–Hochberg FDR correction. All statistical tests were two-sided. For all statistical analyses p-value considered significant if < 0.05.

## Results

### Lake Kinneret physico-chemical environmental conditions differ between sampling layers and season

To identify the chemical parameters characteristic of the different sampling layers and seasons (i.e., stratified vs. mixing periods of the water column), we performed a PCA analysis (n=84). For a full description, see the PCA analysis section in the materials and methods section. We narrowed the initial number of indicative environmental parameters to 6: Temperature, DO, SO4^2-^, pH, NH4^+^ and chlorophyll (Fig. 2A, Fig. S2 and Table S11). Based on these 6 parameters, distinct groupings were noted, based on both sampling layer and sampling season (PC1-PC2: 60.86-18.39%; Fig. 2A). The epilimnion layer was characterized by higher temperatures, DO and chlorophyll levels, with moderate basic pH conditions (pH=8.37±0.006). The metalimnion showed slightly lower temperatures, moderate DO and chlorophyll levels and slightly less basic pH (pH=8.04±0.009). SO4^2-^ and NH4^+^ levels were similar between the two. Temperature, DO, chlorophyll, pH (pH=7.64±0.02) and SO4^2-^ were all lower for the hypolimnion. The only parameter, among the 6 indicative parameters discussed above, showing elevated levels in this layer was NH4^+^, which was also characterized by higher variability. In line with this, samples of the hypolimnion portrayed the greatest separation, compared to the other sampling layers (Fig. 2A). In the samples taken during the water mixing period (Mixis); Temperature, DO, chlorophyll, pH and NH4 were typically lower with only SO4, showing elevated levels.

### *Legionella* spp. occurrence increases with depth peaking in the anoxic hypolimnion

*Legionella* spp. absolute abundance showed a very similar pattern of distribution in all three sampled layers – epilimnion, metalimnion and hypolimnion (Fig. 2B). Yet, lower levels were measured at the epilimnion compared to the mostly anoxic hypolimnion (F(2, 60)=3.67, *p=0.031*; Tukey test, *p=0.*0235, Fig. 2B and Table S1). To gain a higher resolution of *Legionella* spp. absolute abundance in the water column, we collected two depth profiles during March 2018 (n=6, mixis period) and June 2018 (n=16, stratified period), and conjointly measured the following environmental parameters: Temperature, DO, chlorophyll, pH, Ptot, SO4^2-^, NH4^+^. Interestingly, both profiles showed a very similar pattern, with abundance increasing with depth up to 15m (March) and 18m (June). From thereon levels remained relatively constant up to 25m in both profiles (Fig. 2C), with some fluctuations in the June profile. Although significant correlations were noted during the March profile between abundance and temperature, DO, chlorophyll and pH (rs=-0.943, *p=0.005*, Table S2); these correlations are nor reliable due to the small sample size and cross correlation between environmental parameters (Fig. 2C, upper panel). As for the June profile, significant correlations were noted for temperature (rs=-0.58, *p=0.022*) and DO (rs=-0.51, *p=0.05*), and on the verge of significance with pH (rs=-0.5, *p=0.057*) (Fig. 2C, lower panel and Table S3). Notably, similar to what was observed in the monthly profiles, the June profile also revealed high *Legionella* absolute abundances at depths characterized by anoxic conditions. These results are unexpected as *Legionella* is considered an aerobic bacterium. Moreover, our results show a significant negative correlation of *Legionella* abundance with DO concentrations.

### Legionella spp. occurrence peaks during and shortly after water mixing

To assess the variations in *Legionella* absolute abundance between the sampling months, we performed an LMM analysis, setting time as a fixed effect, while controlling for variations related to layer (random effect). Although overall absolute abundance did not differ significantly between the mixis and stratified periods (Table S4). Peaks in *Legionella* abundance were noted during the mixis periods and up to two-three months after the mixis period ended (F(29) = 11.934, *p = 7.67e-11*, Tables S5-S8 and Supplementary Data File 1). This included May-July 2017, March and April 2018, March and June 2019 (Fig. 2B lower panel, marked in gray boxes). During subsequent months, lower (fluctuating) levels were observed, which increased in the following mixis period. Our LMM analysis focused on the metalimnion and hypolimnion layers, as the lower levels noted at the epilimnion layer, significantly increased the variability between sampling dates and masked the observed differences.

### Environmental parameters influencing *Legionella* spp. abundance

Our random forest (RF) analysis indicated the main abiotic parameters that influenced *Legionella* absolute abundance were temperature and Ptot, followed by nitrate and chlorophyll levels (n=84, R^2^ = 0.903, OOB = 0.298, RMSE = 0.838, Fig. S3A). Although the model explains approximately 90% of the variance in *Legionella* abundance based on the selected environmental parameters, the Out-Of-Bag (OOB) score was 0.298, indicating moderate generalization ability to unseen data. The model achieved a Root Mean Squared Error (RMSE) of 0.838, reflecting high prediction accuracy. The biotic RF model indicated the main parameter that influenced *Legionella* spp. abundance was dinoflagellate biomass. However, the biotic model did not perform well (n=30, R^2^ = 0.79, OOB = -0.54, RMSE = 1.005, Fig. S3B). While the R^2^ value demonstrated moderate performance, the OBB was -0.54, indicating poor generalization ability from the trained to the unseen-test data and may indicate overfitting of the model. The model achieved a Root Mean Squared Error (RMSE) of 1.005, reflecting high prediction accuracy. These results suggest that while the biotic model may have captured some factors influencing *Legionella* spp. abundance, the low OBB indicate significant improvements are needed to enhance the model’s generalization capability. To complement the RF analysis, we also performed linear regression between *Legionella* counts [cells / ml] to selected biotic and abiotic parameters. Of the tested parameters, temperature (rs=-0.41, *p<0.001*), chlorophyll (rs=-0.423, *p<0.001*) and pH (rs=-0.34, *p=0.0014*) were significant (Table S12), suggesting the relationships with the other parameters were not linear. We did not perform multiple regression due to cross correlation between variables. Taken together, these results suggest temperature, Ptot, nitrate, chlorophyll, pH and potentially dinoflagellate biomass were the main environmental parameters that influenced *Legionella* abundance. Our depth profiles also showed a negative correlation with DO. This might be due to the higher depth resolution observed in these profiles compared to the RF analysis, which only included three depths per sampling date.

### *Legionella* spp. microbial community composition shifts are related to sampling month and DO concentrations

The composition of the *Legionella* community was determined using genus- targeted amplicon sequencing. We identified *Legionella* ASVs in all samples (n=28, Supplementary Data File 3). A total of 79 *Legionella* ASVs were identified, most were shared between the epilimnion, metalimnion, hypolimnion and mixis samples (Fig. 3A, left panel). About 20% were layer specific (epilimnion, 10.1% and metalimnion 8.9%). The hypolimnion layer and mixis period samples, did not harbor specific ASV. Conversely, approximately 50% of the ASVs were month specific (Fig. 3A, right panel). Furthermore, a pattern indicating a shift in *Legionella* community composition, that is based on sampling month and DO concentrations, but not depth, was found - as is exemplified in the ASVs heatmaps (Figs. 3B-3C). Interestingly, during the mixis period ASVs are shared between the months, with no such apparent shift. While some of the ASVs present during mixis were also present during other months of the year, a gradual change was noted (Fig. 3B). Shared ASVs were also observed when samples were ordered based on DO concentrations. A total of 45 ASVs were identified in anoxic samples. Yet, these were not unique to anoxic samples and are thus not indicative of anoxic conditions. That said, several ASVs were unique to hypoxic conditions (15m and 25m, Apr 2018 and Jun 2019, <2 mg/L; 6a309, e90ea, d1064 and 27acd). In addition, several ASVs that were found under anoxic conditions, were not present at the oxic layers of the corresponding sampling date (i.e., 15m and 25m, June 2018 - 1b5c3, a3ca6, f576f, 6c215, 60a62, 104d7, 7930d, 9e5c2, 83bd3; 25m Aug 2018 -

**Figure 3.**
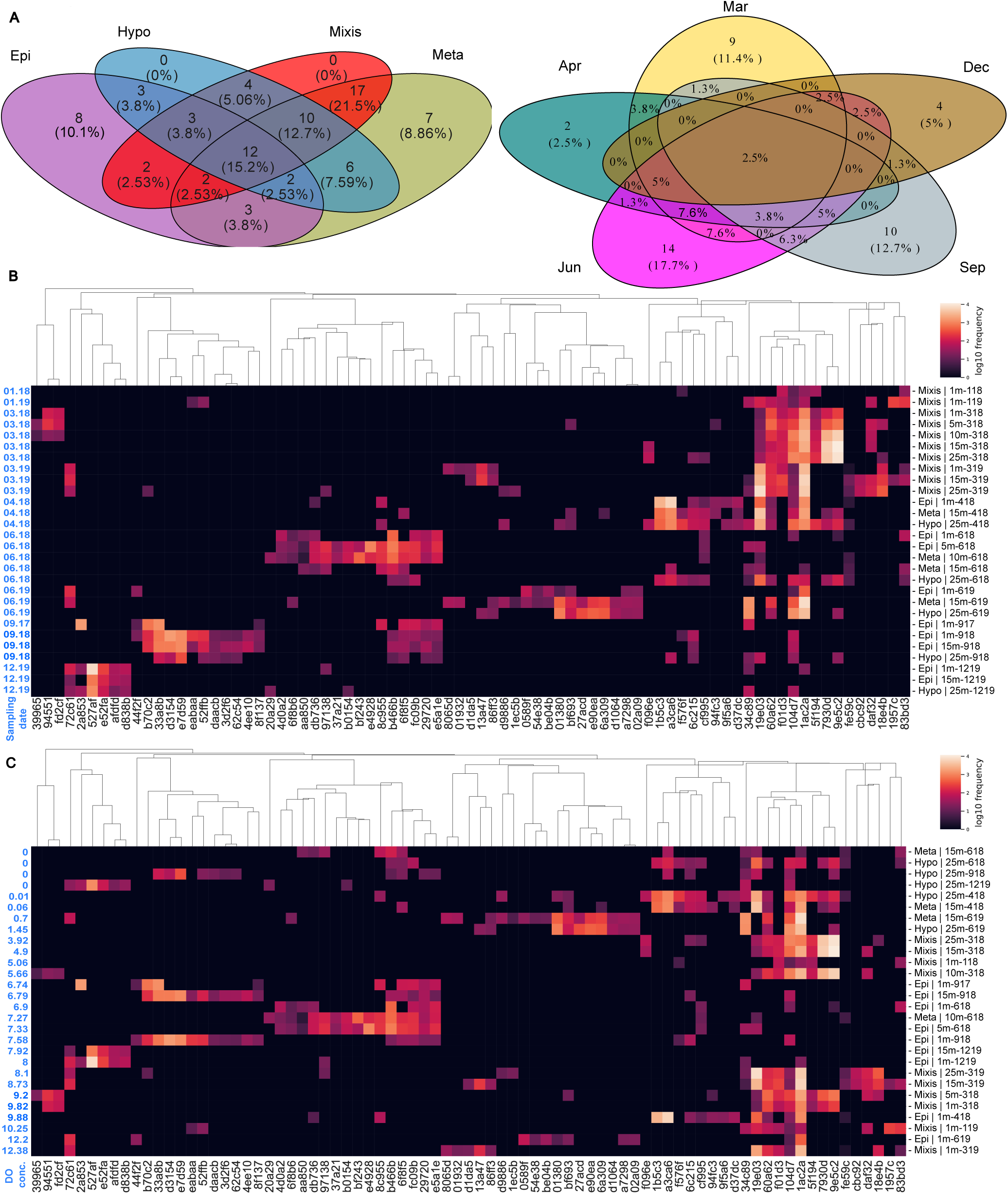
***Legionella* spp. community composition pattern shifts based on sampling month and DO concentrations.** Next Generation Sequencing (NGS, n=28) was performed using *Legionella* spp. specific 16S primers. Samples included water column stratification layers: Epilimnion (Epi=purple, n=9), metalimnion (meta=pale-blue, n=4) hypolimnion (Hypo=mustered-green, n=5), and samples from the complete water column mixing period (mixis=maroon, n=10). **(A)** Venn diagram of *Legionella* ASVs (minimum of 20 reads) separated according to sampling layer (left panel) and sampling month (right panel). Sampling months included March (Mar = light-yellow, n = 8), April (Apr = teal, n = 3), June (Jun = pink, n = 8), September (Sep = gray, n = 3) and December (Dec = brown, n = 3). Three January samples were are not included in the Venn diagram to simplify the plot. (**B+C**) Visualizations presented as relative abundance heatmaps of *Legionella* ASVs (minimum 20 reads). Rows represent samples, ordered according to **(B)** sampling date (and depth - within each date), and **(C)** Dissolved Oxygen (DO) concentrations. DO concentrations are presented to the left of the heatmap. Columns represent ASVs grouped according to Hierarchical-clustering, based on their distribution among the samples. The heatmap color scale represents relative abundance of the ASVs within each sample, from low (blue to black), to high (while-yellow-orange), according to the color bar. ASVs identity is represented by alphanumeric codes at the bottom of each heatmap. For the full description of each ASV, see Supplementary Data 3.

a3ca6, 34c89, 19e03, 9e5c2; and December 2019 – 20a29, b0154, 104d7). A BLAST search performed on the sequences of all these ASVs, identified in oxygen limiting conditions, validated all are indeed related to uncultured or unidentified *Legionella* (Supplementary Data File 8).

A constant and significant increase in alpha diversity was noted with a progression from one month to the other, starting in December (adjacent and prior to the mixis period) through September, for both Shannon’s entropy index (H4=13.96, *p=0.007*) and Faith’s PD indices (H4=11.87, *p=0.018*; Fig. 4A). Post Hoc analysis indicated significantly lower Shannon’s entropy in December compared to both June and September and lower Faith’s PD in March compared to June and September. Showing a similar trend, Pielou’s evenness index did not differ significantly (Fig. 4A). All Kruskal–Wallis and Post Hoc via Wilcox tests with BH corrections are presented in Tables S13-S14. The only significant difference in alpha diversity between the different sampling depth layers was observed in Faith’s PD index (H3=9.64, *p=0.021*; Fig. S5, Tables S13, S15), with lower diversity during mixis compared to the Hypolimnion. Beta diversity showed a very clear separation between sampling months in both Jaccard index (PERMANOVA, F5,21=5.18, *p=0.001*, Fig. 4B left panel) and less profoundly in the Bray Curtis index (PERMANOVA, F5,21=5.14, *p=0.001*, Fig. 4B right panel). In contrast, beta diversity of the sampling depth layers, indicated a difference only in the Jaccard index between mixis and epilimnion (PERMANOVA, F4,21=1.78, *p=0.014*, Fig. S6). PERMANOVA, PERMDISP and

**Figure 4.**
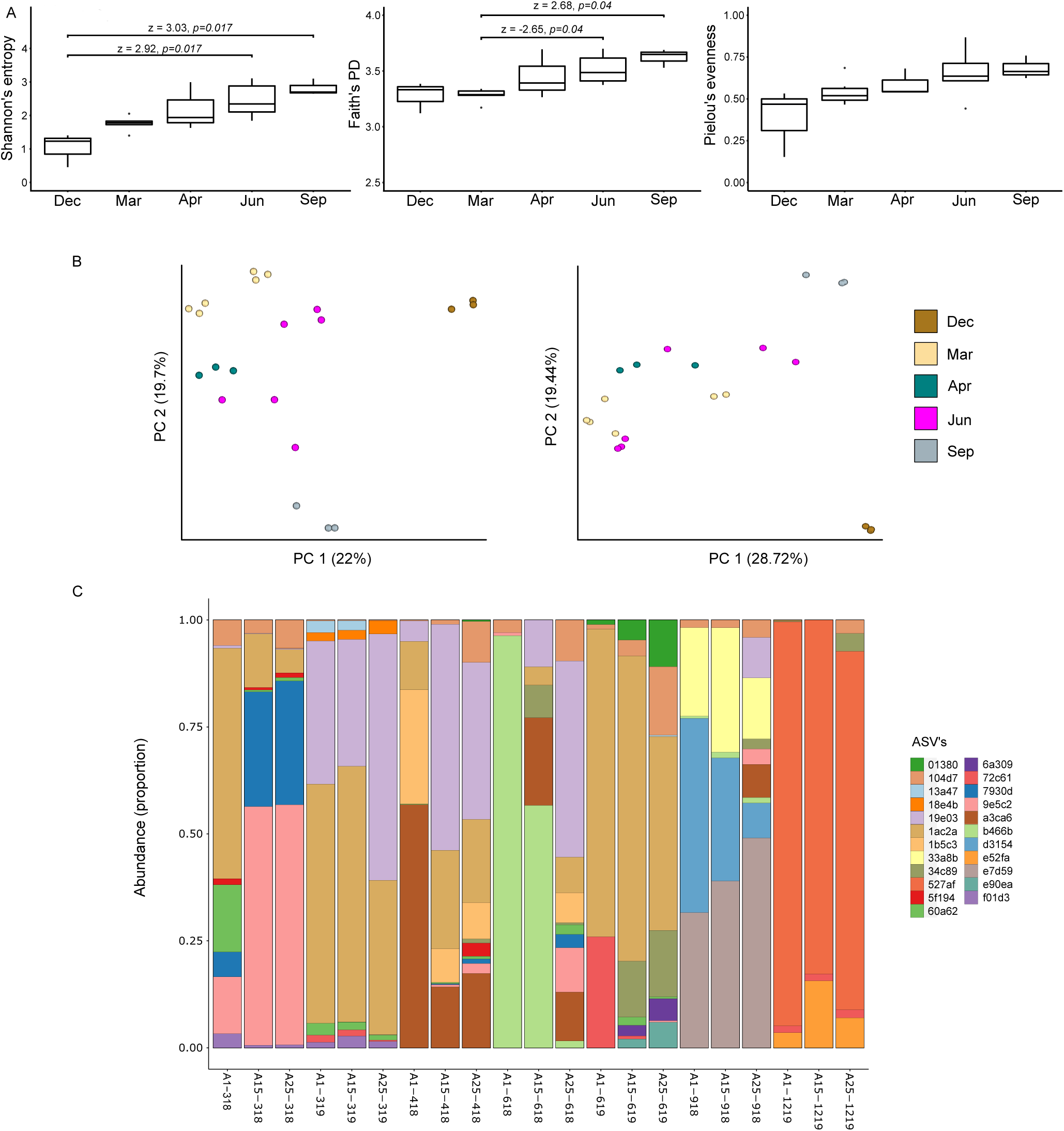
***Legionella* ASV diversity shifts according to month, with dominant- alternating ASVs.** Next Generation Sequencing (NGS, n=21) was performed using *Legionella* spp. specific 16S primers. ASVs for the diversity metrices analysis were pre- filtered to a minimum frequency of 20 reads, across a minimum of two samples. To study *Legionella* diversity according to sampling month, while maintaining a relatively balanced study design, we included in the analysis only the months that were sampled at all three depths. In addition, we performed a separate diversity analysis based on the water column sampling layer (Figs. S5-S6). Samples included in the analysis were: March (Mar= light-yellow, n = 6), April (Apr = teal, n = 3), June (Jun = pink, n = 6), September (Sep= gray, n = 3) and December (Dec = brown, n = 3). **(A)** Alpha diversity of *Legionella spp.* according to sampling month. Shannon’s entropy (left panel), Faith’s PD (middle panel) and Pielou’s evenness (right panel) are presented. Kruskal-Wallis test results and post hoc via Wilcox tests are presented in Tables S13-S14. **(B)** Principal coordinate analysis plot depicting differences in the *Legionella spp.* microbial community according to sampling month, based on the beta diversity indices Jaccard (left panel) and Bray–Curtis (right panel). Statistical significance was tested via PERMANOVA and PERMDISP analyses, Number of permutations = 999 (Supplementary Tables S16-S17). **(C)** Barplot visualization of abundant *Legionella* ASVs relative abundance (minimum frequency of 500 reads, across a minimum of two samples). Samples were ordered by sampling month and depth.

Post Hoc results are presented in Tables S16-S17). We conducted PERMDISP tests, to help assess whether the differences observed in beta diversity among sampling months are due to differences in the average composition (centroid) or due to differences in the variability (dispersion) within each group. While both PERMDISP and PERMANOVA were for the most part significant (Table S16), the clustering pattern suggests that the differences are due to differences between the sampling month. The effects of environmental conditions on beta diversity were analyzed via the ADONIS test (Table S18) with the following variables: Temperature, Cl, DO, pH, Ptot, NH4, month and layer). While all variables were significant for both Jaccard and Bray Curtis, aside from the layer variable in the Bray Curtis index. The variable explaining most of the variability was month for both Jaccard and Bray Curtis (R^2^=0.283, R^2^=0.323, respectively). Temperature (R^2^=0.116, R^2^=0.125, respectively), Cl (R^2^=0.142, R^2^=0.138, respectively) and pH (R^2^=0.084, R^2^=0.112, respectively) followed by DO (R^2^=0.087, R^2^=0.044, respectively). Focusing on the more abundant ASVs (minimum 500 reads, in at least 2 samples), we identified the occurrence of dominant *Legionella* ASVs in each of the sampling months. Most months were characterized by these dominant ASVs, which seemed for the most part to be month specific (Fig. 4C). Few exceptions were noted; mainly ASV 1ac2a (Blast search: Uncultured *L. bozemanae*, 98.4% identity), which was present in most sampling months in high relative abundance (omitting September) and ASV 19e03 (Blast search: *L. adelaidensis*, 97.86% identity), which was observed in March, April, June and September.

Taken together *Legionella* spp. microbial community composition and the diversity analysis, reflected the results obtained in the qPCR analysis. The main differences were temporal (month), rather than spatial (layer), with environmental parameters such as temperature, DO, Cl and pH explaining the variations in *Legionella* community composition.

### Phylogenetic analysis

To test whether the monthly shifts in *Legionella* community composition were associated with phylogenetically related *Legionella* species, we constructed phylogenetic trees with known *Legionella* species (Supplementary Data File 4). When all ASVs were included in the phylogenetic analysis (filtered minimum 20 reads, in at least 2 samples), no phylogenetic clades associated with the sampling month or layer were observed (Fig. S7-S8). When examining only the most abundant *Legionella* (>15%) we identified a phylogenetic clade consisting of exclusively June related genotypes, closely related to the *L. israelensis* species (72c61, b466b and 8c955, Fig. 5). Another small phylogenetic clade was identified featuring two genotypes, both from March (7930d and 9e5c, Fig. 5). The two dominant ASVs identified in multiple months: 19e03 (Blast search: *L. adelaidensis*, 97.86% identity), and 1ac2a (Blast search: Uncultured *L. bozemanae*, 98.4% identity), were distantly related to *L. adelaidensis* and *L. steigerwaltii* on the tree (respectively). Of the total ASVs (20 reads filtered, 79 features) only a few were closely related to known *Legionella* species on the tree (Figs. S7-S8). These include: *L. feeleii* (ASV d838b), *L. rubrilucens* (37a21), *L. waltersii* (2a853), *L. worsleiensis* (6f8b6), *L. steigerwaltii* (02a09 and e7d59). Notably, ASV 44f2f was classified as *L. pneumophila* (99.2% identity) at the epilimnion only, during June and September. Interestingly, of the four *Legionella* ASVs unique to hypoxic conditions, 6a309 and e90ea clustered closely together on the same phylogenetic clade. In agreement with the aforementioned BLAST results, all ASVs identified in the oxygen limiting conditions were not related to known *Legionella* species.

**Figure 5.**
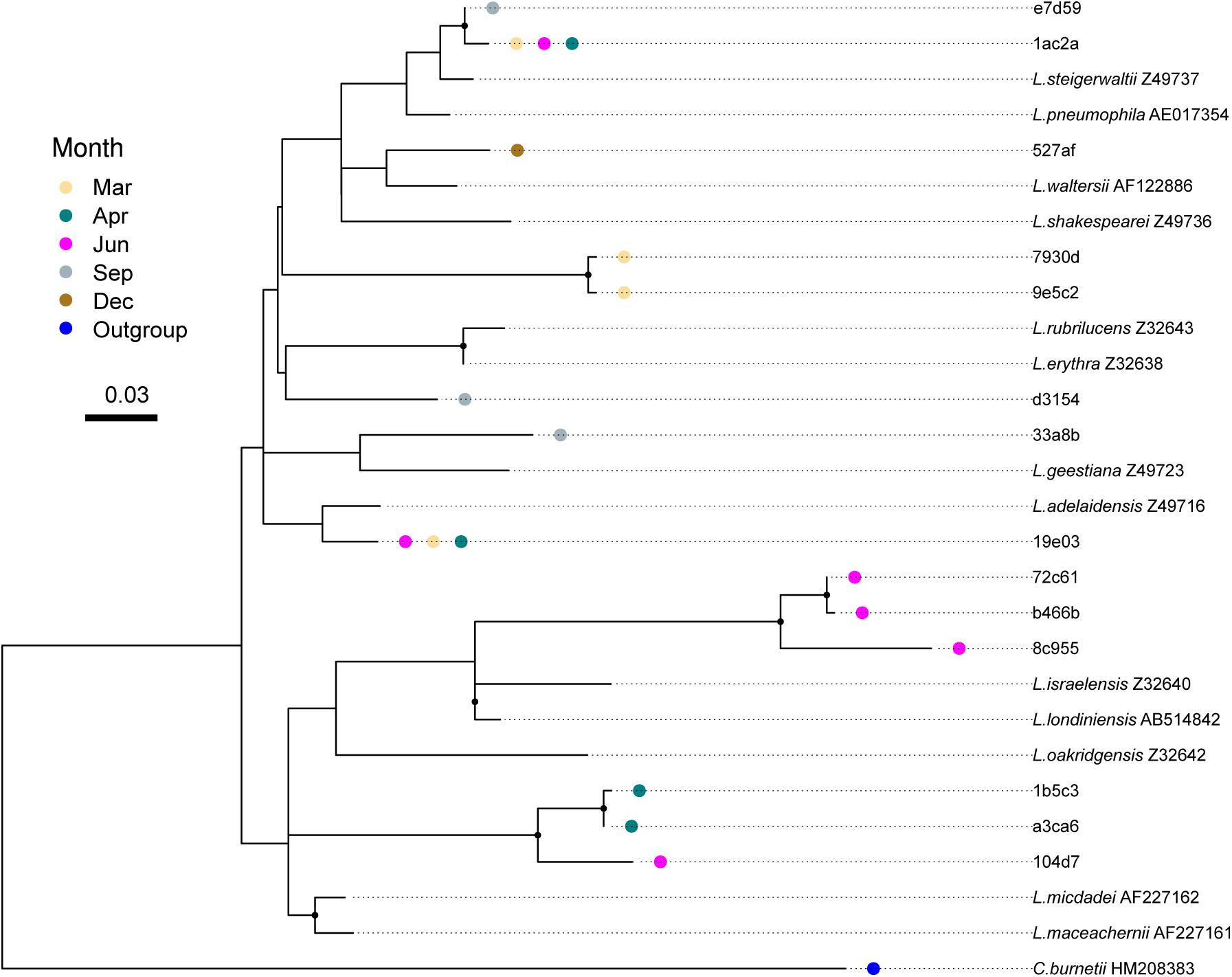
**Phylogenetic analysis of *Legionella* ASVs shows a monthly based grouping**. The phylogenetic tree included dominant ASVs, raw data was filtered to included ASVs with a relative abundance >15%. The tree was constructed alongside reference bacteria with corresponding NCBI accession-numbers (Supplementary Data 4). For readability only closely related *Legionella* reference bacteria were included in the final tree construction. Month and Leyer based trees of all ASVs (filter 20 reads, in at least 2 samples) are also The Maximum-Likelihood method and Hasegawa-Kishino-Yano (HKY) model were used to infare the tree evolutionary history. We included as an outgroup *Coxiella burnetii*. The tree with the highest log likelihood is presented (-3895.04). A discrete Gamma distribution was used to model evolutionary rate differences among sites (5 categories (+G, parameter = 0.2031)). The rate variation model allowed for some sites to be evolutionarily invariable ([+I], 30.29% sites). The tree is drawn to scale, with branch lengths measured in the number of substitutions per site. Branches with bootstrap values above 50% are demoted by a black dot. The analysis involved 75 nucleotide sequences. There was a total of 379 positions in the final dataset. Evolutionary analyses were conducted in MEGA11.

### Microbial community composition of potential *Legionella* hosts

The ability of *Legionella* spp. to replicate within various hosts, such as Amoebozoa, Ciliophora, and Percolozoa, is believed to contribute to their survival, persistence and dispersal in the environment [28–30,33]. To study the occurrence of *Legionella* spp. with potential protozoan hosts, we conducted an 18S rRNA amplicon-based sequencing, targeting protozoa and filtered the resulting ASVs to include representatives related to groups of known *Legionella* hosts: Amoebozoa, Ciliophora, and Percolozoa. In addition, we included Dinoflagellates, as they were found to potentially influence *Legionella* absolute abundance in the Random Forest algorithm. A total of 163 potential host ASVs were identified. Similar to the identified *Legionella* ASVs, our community composition analysis indicated approximately 23% of the these were layer specific, predominantly in the epilimnion (8.5%). Contrary, the hypolimnion and mixis layers, which did not harbor specific *Legionella* ASVs, showed specific potential host ASVs (4.9% and 5.5%, respectively). On the other hand, approximately 23.3% of the ASVs were month specific (compared to 50% of *Legionella* ASVs) (Fig. 6A). While differences in the potential *Legionella* host community were observed between the mixis and stratified periods, these were significantly reduced compared to the shift observed in the *Legionella* community as can be seen in the ASVs heatmaps of both (Figs. 3B, 6B). The majority of ASVs were of Ciliophora (62%), Dinoflagellata (33.8%) and only 4.2% Amoebozoa (Fig. 7A upper panel and Supplementary Data File 6). Differences were also noted when samples were ordered according to DO concentrations, with a group of genotypes appearing almost exclusively in samples from anoxic conditions (Fig. 6C). Amoebozoa were only identified at low relative abundance (4.2%), in agreement with the low relative abundance observed in water from natural spring habitats surrounding the Sea of Galilee [70]. Most months were characterized by the presence of dominant ASVs (Fig. 7A, bottom panel and Supplementary Data File 6) classified as both Dinoflagellates: 3ed59 (Blast search: *Peridinium polonicum*, 98.4% identity), 2c647 (Blast search: *Parvodinium (Parvodinium) cunningtonii*, 100% identity), 39fc1 (Blast search: *Peridinium cinctum*, 98.4% identity), bfe9e (Blast search: *Ceratium hirundinella*, 100% identity) and Ciliophora: bd959 (Blast search: *Mesodinium pulex*, 100% identity), cab72 (Blast search: *Pelagostrombidium fallax*, 100% identity), 48d60 (Blast search: *Choreotrichia* sp., 100% identity) daff9 (Blast search: *Coleps hirtus*, 100% identity), 70296 (Blast search: *Choreotrichia* sp., 100% identity), 80ce6 (Blast search: *Halteria* sp., 100% identity) and 1b115 (Blast search: *Tintinnidium fluviatile*, 100% identity). The only difference in alpha diversity was lower Faith’s PD (H3=12.64, *p=0.005*; Fig. S10 and Table S19), in the mixis period compared to the metalimnion (*p=0.002*) and hypolimnion (*p=0.033*, Table S20). No difference was found between different sampling months (Fig. 7B and Table S19). Analysis of beta diversity showed differences in the potential host community structure by sampling month, based on both Jaccard and Bray Curtis indices (PERMANOVA, F5,21=4.523, *p=0.001* and F5,21=3.45, *p=0.001*, respectively; Fig. 7C and Table S21). During mixis, significant differences were found compared to the three sampling layers (i.e., stratified periods) (PERMANOVA, F4,21=2.985, *p=0.001* and F4,21=2.435, *p=0.001*, respectively; Fig. S11 and Table S21).

**Figure 6.**
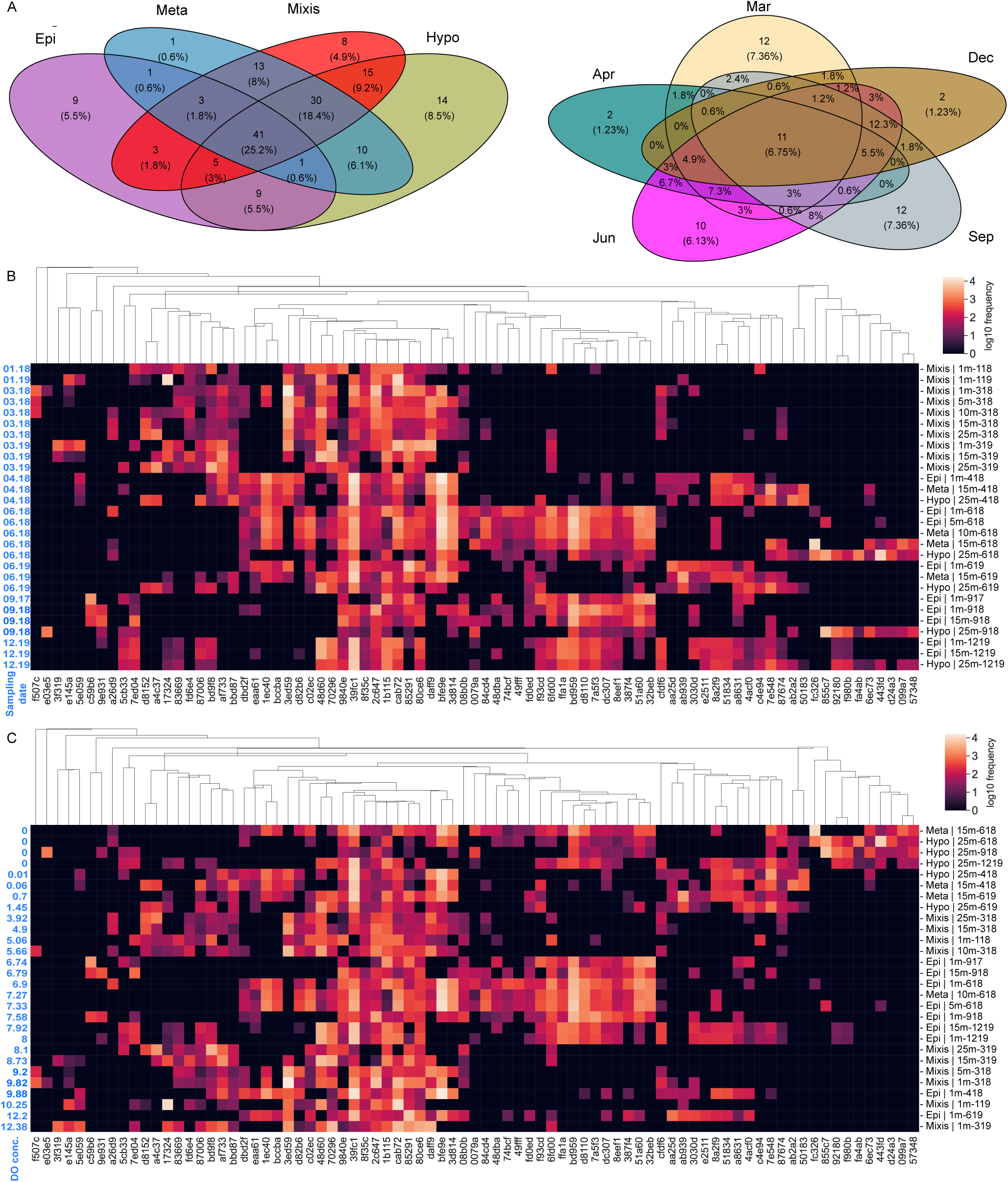
Alterations in community structure and composition of potential *Legionella* hosts between mixis to stratified periods, and according to DO concentrations. Next Generation Sequencing (NGS, n=28) was performed using universal protists primers, designed for the V9 region of the 18S rRNA gene. Raw data were pre-filtered to retain Amoebozoa, Cliophora (Ciliates), Percolozoa (Excavata) and Dinophyta (Dinoflagellates). We included in the analyses water column samples from stratified layers: Epilimnion (Epi=purple, n=9), metalimnion (meta=pale-blue, n=4) hypolimnion (Hypo=mustered-green, n=5), and samples from the complete water column mixing period (mixis=maroon, n=10). **(A)** Venn diagram of *Legionella* hosts ASVs were prefiltered to retain ASVs with a minimum of 20 reads, in at least 2 samples. Samples were separated according to layer (left panel) and sampling month (right panel). Sampling months included March (Mar = light-yellow, n = 8), April (Apr = teal, n = 3), June (Jun = pink, n = 8), September (Sep = gray, n = 3) and December (Dec = brown, n = 3). We omitted January samples to simplify the Venn diagram. (**B+C**) Relative abundance heatmaps of *Legionella* hosts ASVs (filtered 500 reads, in at least 2 samples). Samples, were ordered in rows according to **(B)** month-year (and depth within each combination), and **(C)** Dissolved Oxygen (DO) concentrations (presented to the left of the heatmap). ASVs were grouped in columns according to Hierarchical-clustering, based on distribution among the samples. The heatmap color scale represents ASVs relative abundance, with larger levels corresponding to higher values (while-yellow-orange) and vise-versa (blue to black). ASVs identity is presented by id’s at the bottom of each heatmap. For the full description of each ASV, see Supplementary Data 6.

**Figure 7.**
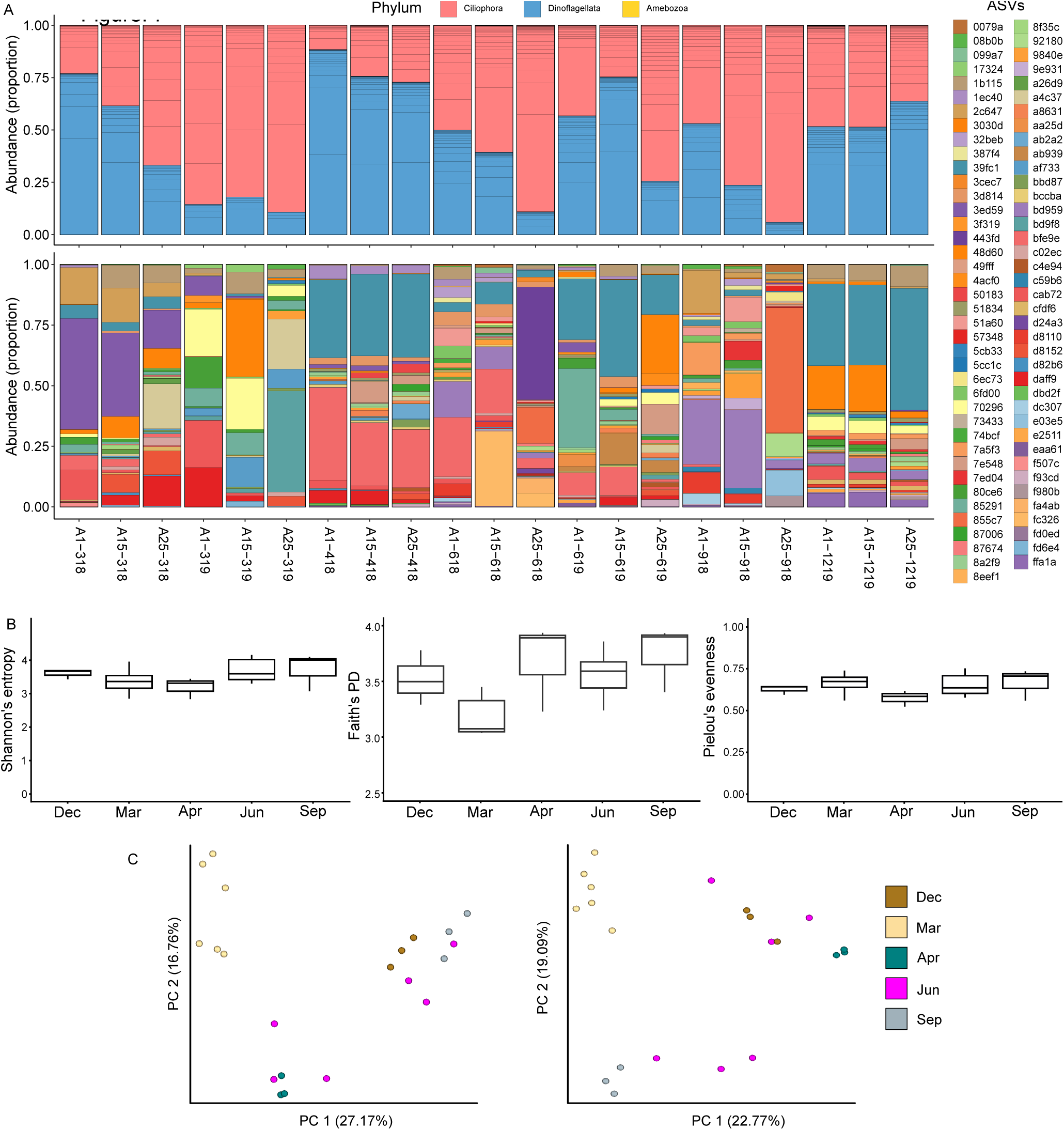
Potential *Legionella* hosts portray variations in beta diversity, with dominant ASVs monthly based alteration. *Legionella* spp. specific 16S primers were used to perform Next Generation Sequencing (NGS, n=21). ASVs for the diversity metrices analysis were pre-filtered to a minimum frequency of 20 reads, across a minimum of two samples. To Maintain a relatively balanced study design, we included in the month-based diversity analysis, only months that were sampled at all three depths. In addition, we performed a separate diversity analysis based on the water column sampling layer (Figs. S10-S11). Samples included in the analysis were: March (Mar = light-yellow, n = 6), April (Apr = teal, n = 3), June (Jun = pink, n = 6), September (Sep = gray, n = 3) and December (Dec = brown, n = 3). **(A)** Shannon’s entropy (left panel), Faith’s PD (middle panel) and Pielou’s evenness (right panel) were included in the alpha diversity analysis. Kruskal-Wallis test results and post hoc via Wilcox tests are presented in Tables S19-S20. **(B)** Beta diversity of *Legionella spp.* potential hosts was performed according to sampling month. Jaccard (left panel) and Bray–Curtis (right panel) indices are presented. Beta diversity statistical significance was done via the PERMANOVA and PERMDISP analyses, Number of permutations = 999 (Supplementary Tables S21-S22). **(C)** Relative abundance barplot visualization was performed for abundant potential *Legionella* ASVs hosts. ASVs at the phylum level (top panel, minimun 20 reads) and at the ASV level (bottom panels, minimum 500 reads) are presented. Samples were ordered by sampling month and depth.

### Legionella co-occurrence patterns with potential hosts differ between oxic and anoxic environments

To account for the identification of *Legionella* ASVs in anoxic environments, we wanted to test for possible interactions with potential *Legionella* hosts. Such interactions have been reported, sustaining *Legionella* under harsh conditions, while creating microenvironments beneficial for their survival (Abdel-Nour et al., 2013; Boamah et al., 2017). We performed separate cross-domain co- occurrence network analyses, for oxic and anoxic environments. Based on the NGS amplicon sequencing of both groups, utilizing the SPIEC-EASI package. Our analysis indicated that *Legionella* ASVs from water samples of anoxic conditions (n=6) portrayed similar taxonomy-based interactions, with both Dinoflagellates (14 edges) and Ciliophora (17 edges, Fig. 8A). In comparison, the co-occurrence network of *Legionella* ASVs from oxic samples (n=13), interacted more with Ciliophora (68 edges) compared to Dinoflagellates (37 edges, Fig. 8B). Several interactions were also observed with Amoebozoa (4 edges).

**Figure 8.**
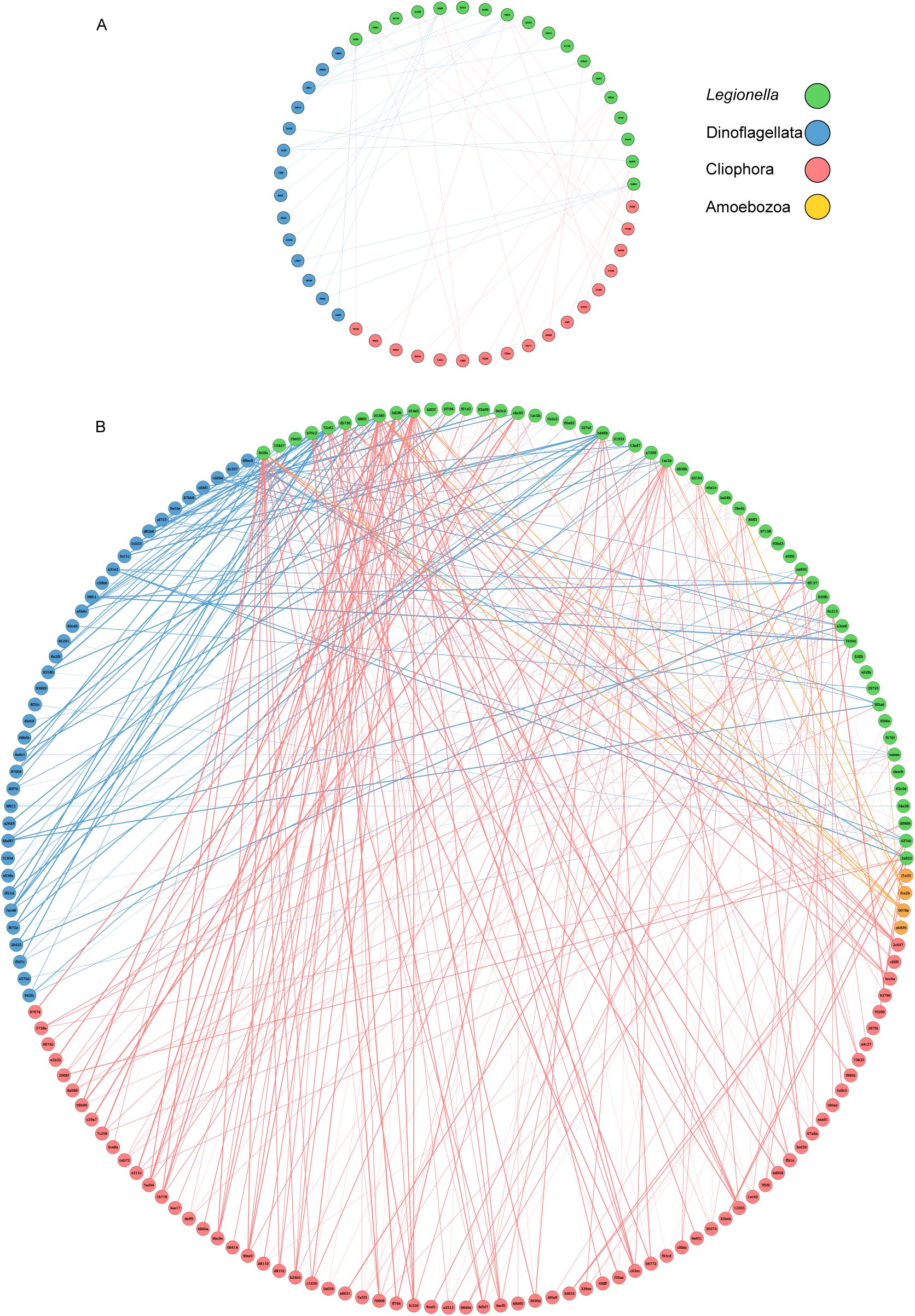
Co-occurrence of *Legionella* ASVs with potential protozoan hosts, in anoxic and oxic environment reveals distinct interactions. SPIEC-EASI cross- domain co-occurrence networks of *Legionella* ASVs and potential-protozoan hosts, associated with samples associated with **(A)** anoxic (n=6) and **(B)** oxic (n=13) conditions. The networks were constructed using Next Generation Sequencing (NGS) with *Legionella*-specific 16S primers, pre-filtered to retain only *Legionella* ASVs. NGS targeting potential protozoa hosts was carried out using 18S rRNA primers targeting the V9 region. Raw data were pre-filtered to retain Amoebozoa, Cliophora and Percolozoa and Dinoflagellate ASVs. ASVs were filtered for a minimum frequency of 20 reads, in a minimum of 2 samples. ASVs are represented by the last 5 letters-number code. For the full-length code and corresponding sequences, see Supplementary Data 3 and 5. Edges with low weights (>0.01) and unconnected nodes were removed. Only positive associations between ASV nodes are presented. Edge width corresponds to associations strength between ASVs.

We identified, for both separately composed networks, co-occurrence with Dinoflagellates of the class Dinophyceae only (Orders Gonyaulacales, Gymnodiniphycidae, Peridiniales and Peridiniphycidae), albeit these were much more frequent in the oxic samples (Supplementary Data 7). A similar pattern was also noted for Ciliates. Interactions were observed with the class Intramacronucleata (Orders Conthreep, Litostomatea and Spirotrichea). For the oxic samples we also observed the class Postciliodesmatophora (Order Heterotrichea). Thus, while the taxonomic nature of interactions was similar, *Legionella* ASVs from oxic conditions tended to co-occur more frequently with Ciliates. Finally, *Legionella* ASVs from oxic samples, tended to interact with more potential host ASVs, compared to those from anoxic samples (e.g., fe59c, db736, 01380, d1da5, b466b, 1ac2a).

## Discussion

The aim of this study was to uncover the forces shaping *Legionella* ecology in the natural environment of a freshwater stratified lake. To this end, we explored *Legionella* spp. absolute abundance, diversity and microbial community dynamics in Lake Kinneret, a sub-tropical, monomictic, freshwater lake. Our results reveal significant seasonal shifts in *Legionella* spp. occurrence and diversity, define the network of interactions with potential hosts and provide evidence for the persistence of *Legionella* genotypes in oxygen deprived freshwater environments. This study challenges the conventional view of *Legionella* as a strictly aerobic pathogen, suggesting that some *Legionella* species have evolved mechanisms to survive conditions of oxygen limitation.

The data collection spanned 2.5 years, and included both biotic and abiotic parameters. Our results reveal seasonal variation in *Legionella* spp. abundance, with higher levels from winter to early spring, coinciding with the later stages of the mixis period when vertical mixing of the water column occurs in the lake. Higher levels were also observed in the hypolimnion, compared to the epilimnion. This result was unexpected as the hypolimnion was anoxic for much of the relevant sampling period, while *Legionella* are traditionally considered aerobic bacteria [94]. Reports of *Legionella* in anoxic environments are usually overlooked or regarded as artifacts. For example, in the identification of *Legionella* in anoxic water collected from groundwater treatment plants, the authors attributed this to “introduction of (trace amounts of) oxygen at the well head”, as “All cultured *Legionella* species need molecular oxygen for multiplication” [13]. We hypothesize that some of the identified *Legionella* species may exhibit a facultative intracellular lifestyle, residing within protozoan hosts, as certain protozoa are known to survive in low-oxygen environments [95]. This intracellular lifestyle can potentially allow *Legionella* to persist in anaerobic conditions indirectly. It is also possible that vertical movement of water masses, may cause occasional mixing events that can transport aerobic *Legionella* cells from upper aerobic to lower anaerobic layers, or alternatively to ventilate deeper water layers [96]. While not a viable option for long term survival, it is possible that frequent mixing events of this nature, may introduce viable *Legionella* cells to anaerobic depths. Under this scenario, survival and persistence is not possible. One limitation of the qPCR methodology used to detect *Legionella* is the inability to differentiate between live and dead bacteria. Therefore, some of the detected DNA might represent dead bacteria that sank to the lower layer of the water column. Indeed, anoxic conditions have been shown to facilitate eDNA preservation [97,98]. Nonetheless, using amplicon sequencing we identified a few unique *Legionella* genotypes present only in the hypoxic zone of the lake, suggesting that the qPCR signal represents, at least in part, live *Legionella* genotypes found in oxygen deprived layers of the lake. It is conceivable that specialized *Legionella* genotypes have evolved under hypoxic and anoxic conditions. These may potentially switch between aerobic and anaerobic metabolic pathways. Under anoxic environments, specialized genotypes may potentially utilize alternative electron acceptors, such as nitrates or sulfates, to sustain their energy production. The occurrence of *Legionella* species in natural hypoxic and anoxic environments needs to be further examined in future enrichment, isolation or transcriptomics experiments in order to gain a better understanding of possible mechanisms facilitating *Legionella* survival under oxygen limitation.

To study *Legionella* microbial community composition, we used genus specific primers and identified 79 *Legionella* ASVs. Our analysis found significant temporal rather than spatial variations in diversity and community composition, reflected in monthly shifts of highly abundant *Legionella* ASVs, and dynamic changes in beta diversity. An accompanying phylogenetic analysis revealed that most ASVs were not closely related to known *Legionella* species. High *Legionella* diversity has also been reported in an ecological survey of *Legionella* in natural springs [70], in the Tech River [57], in Antarctic natural and man- made environments [54] and in two polar lakes of the Antarctic Peninsula [10]. Overall, these findings, together with the results of the current study indicate that *Legionella* diversity is significantly underestimated.

The relationship between *Legionella* and environmental protists has a profound impact on the evolution of *Legionella* as well as on the population dynamics of protists [99,100]. To examined the interaction between *Legionella sp.* And potential *Legionella* hosts in a natural freshwater environment we utilized 18S amplicon sequencing together with co-occurrence network analysis. Our results demonstrate that the vast majority of potential host ASVs were of Ciliophora and Dinoflagellates, with only 4.2% of the ASVs identified as Amoebozoa. Lower Amoebozoa levels were also observed in water samples collected from freshwater and saline springs surrounding Lake Kinneret [70], as well as in water samples from the Puzih River in Taiwan [101], demonstrating high *Legionella* levels, and low overall amoebae detection rate. Collectively, these results suggest that Ciliates and potentially other organisms may play a major role in *Legionella* proliferation in the environment, and may represent overlooked contributors to *Legionella* evolution.

Interestingly, our co-occurrence network analysis indicated frequent interactions between dinoflagellates and *Legionella sp.* Notably, PCA, random- forest and linear correlation analyses, incorporating various biotic and physico- chemical parameters, also pointed at dinoflagellate biomass as a key environmental factor influencing *Legionella* abundance. This influence was observed alongside other parameters, including temperature, Ptot, nitrate, chlorophyll and pH. This association between *Legionella* spp. and dinoflagellates may represent a previously unidentified ecological interaction, potentially mutualistic or commensal in nature, and warrants further investigation.

## In conclusion

*Legionella* species are widely recognized as human pathogens, with most research historically focused on clinical isolates and occurrence in man-made infrastructures, such as potable water systems, cooling towers and hospitals [41,60,102,103]. Thus, the occurrence of *Legionella* in natural environments has been largely overlooked over the years, despite concerns that these natural ecosystems may act as reservoirs and evolutionary ‘training grounds’ for intracellular pathogens such as *Legionella*, and may aid in the future spread of disease to humans [14,60]. The current study addresses this knowledge gap and highlights significant seasonal and temporal shifts in *Legionella* occurrence, community composition and diversity, offering valuable insights into *Legionella* spp. ecology in natural stratified freshwater lake systems. We report the occurrence of *Legionella* under oxygen limited conditions, suggesting novel mechanisms for energy generation, potentially via previously unidentified anaerobic metabolic pathways. The results highlight the complexity of interactions between *Legionella* and the environment, demonstrating how biotic and abiotic factors, including potential environmental hosts shape the abundance and diversity of *Legionella* in natural environments. Our findings provide a comprehensive view of *Legionella* freshwater ecology, offering a potential contribution to the improved management of freshwater systems and a deeper understanding of the evolutionary forces driving *Legionella* pathogenesis.

## Notes

### Competing Interest Statement

The authors have declared no competing interest.

