## Supplementary material for "Unveiling the Ecology of *Legionella* in a Stratified Freshwater Lake: Seasonal Dynamics, Host Interactions, and Persistence Under Oxygen Limited Conditions": Supplamentary information

**The following are included:**

**Supplementary Figs 1-11**

**Supplementary Tables 1-22**

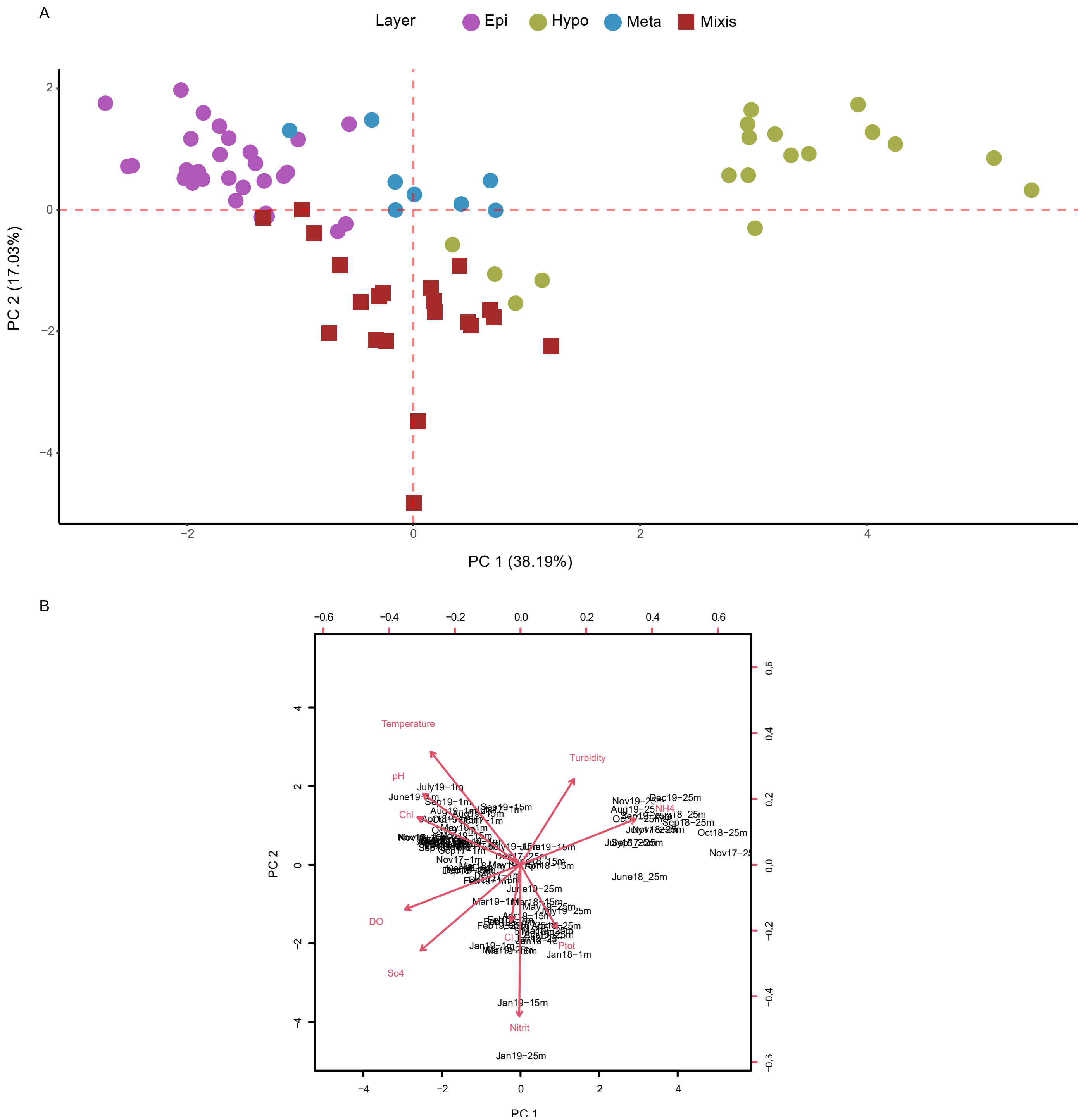


**Figure S1. Initial principal-component analysis (PCA) of various physicochemical characteristics measured at station A.** (A) PCA (n=84) was performed on samples obtained between May 2017 and December 2019 and included 14 environmental parameters (Se materials and methods). After omission of variables presenting strong collinearity (Table S9), the following parameters were included in the initial PCA analysis: Temperature, DO, Cl, SO42-, Turbidity, pH, NH4+, Nitrite, Ptot and Chl (Table S9 and Supplementary data file 2). (B), indicating the importance of tested environmental variables related to PC 1 and PC 2 (Table S10). Based on the loading scores of this initial PCA analysis, a final PCA was performed (Fig. 2C).

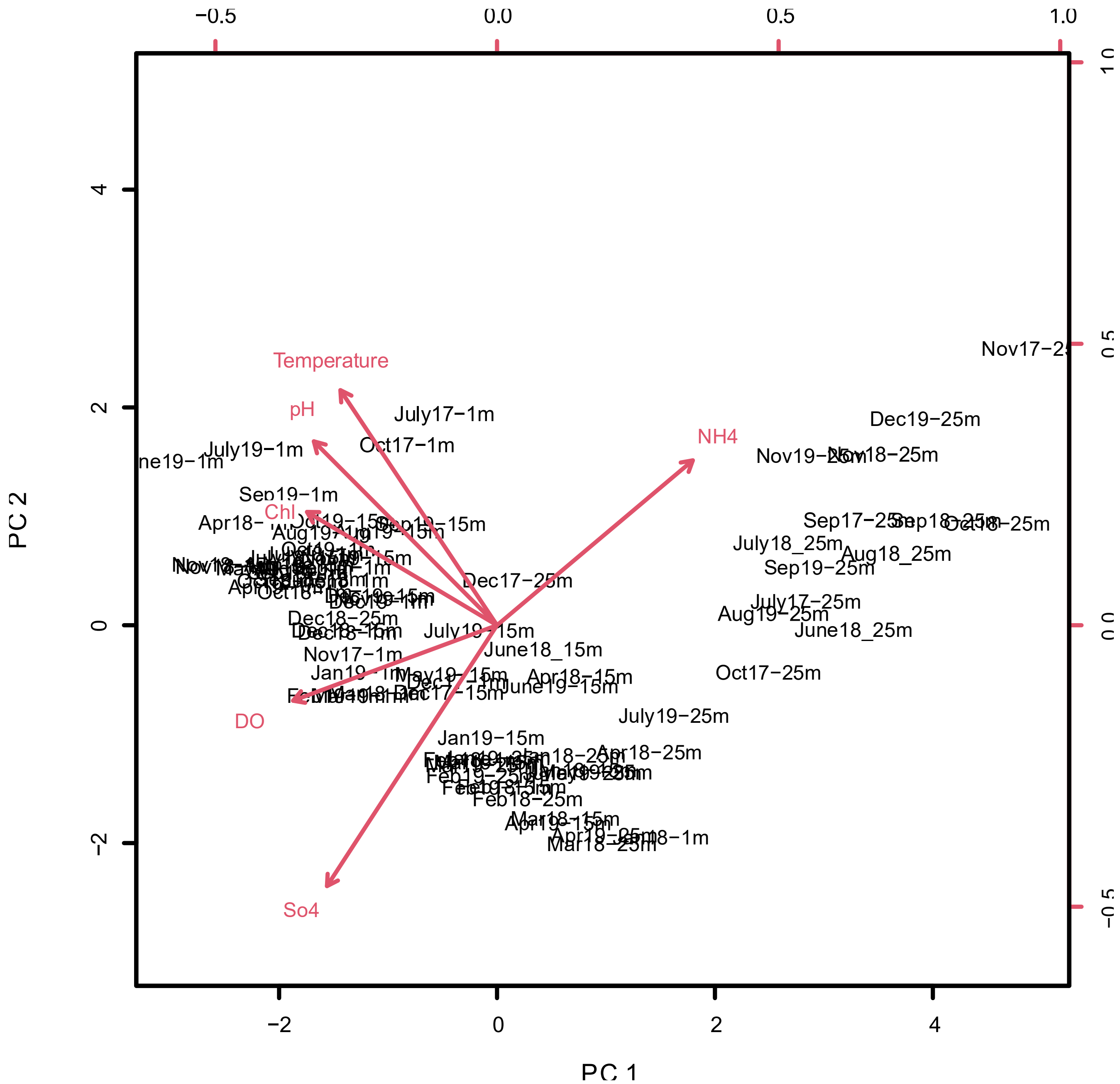


**Figure S2. Loading vectors for the final principal component analysis (PCA), of various physicochemical characteristics measured at station A.** The final PCA was performed on with 6 environmental parameters (Fig. 2C): Temperature, DO, $SO_4^{2-}$, pH, $NH_4^+$ and chlorophyll. Loading vectors indicate the importance of tested environmental variables related to PC 1 and PC 2, based on the loading scores presented in Table S11.

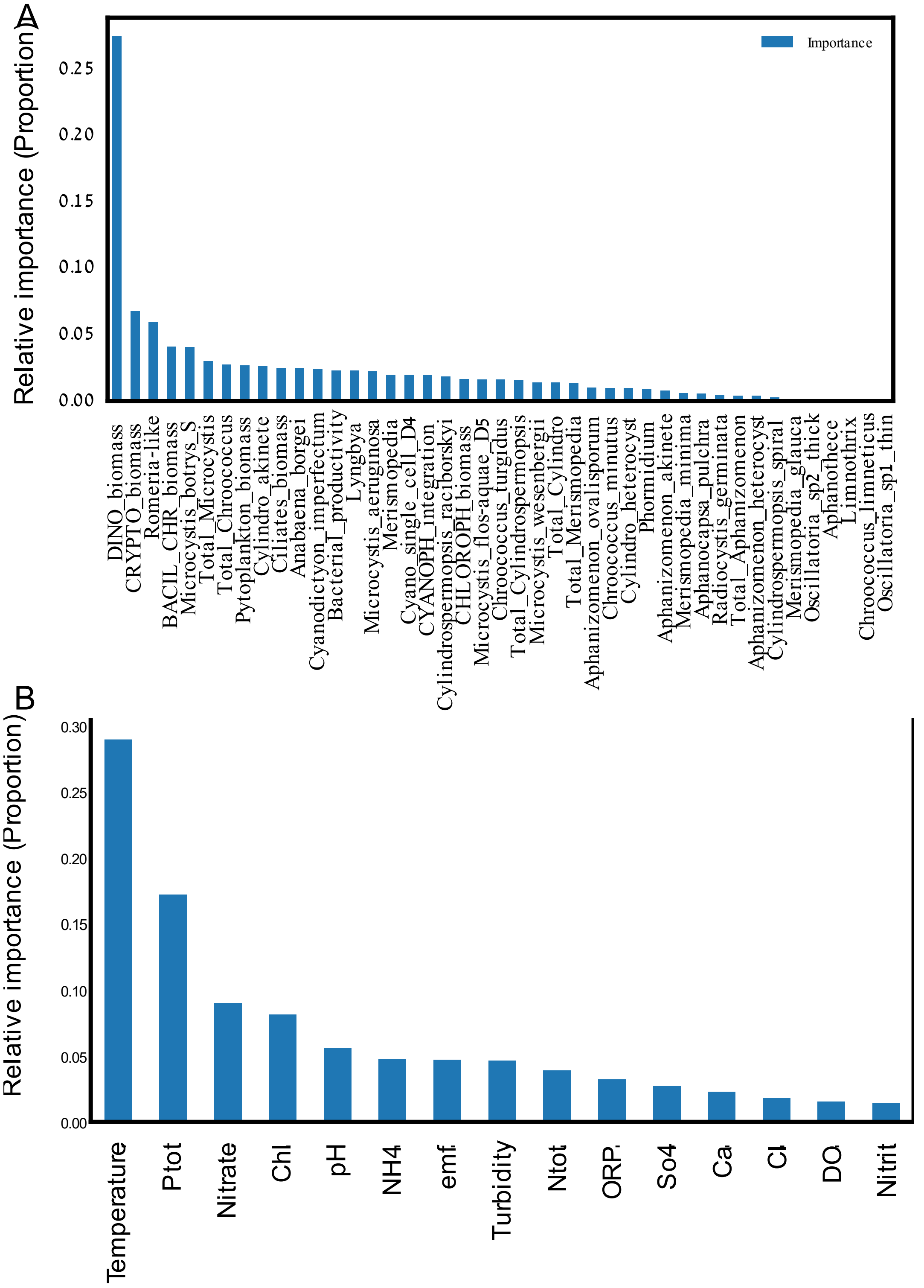


**Figure S3. Relative variable importance in explaining *Legionella* spp. absolute abundance.** Relative variable importance (proportion) was determined for (A) biological and (B) chemical variables, explaining *Legionella* spp. absolute abundance.

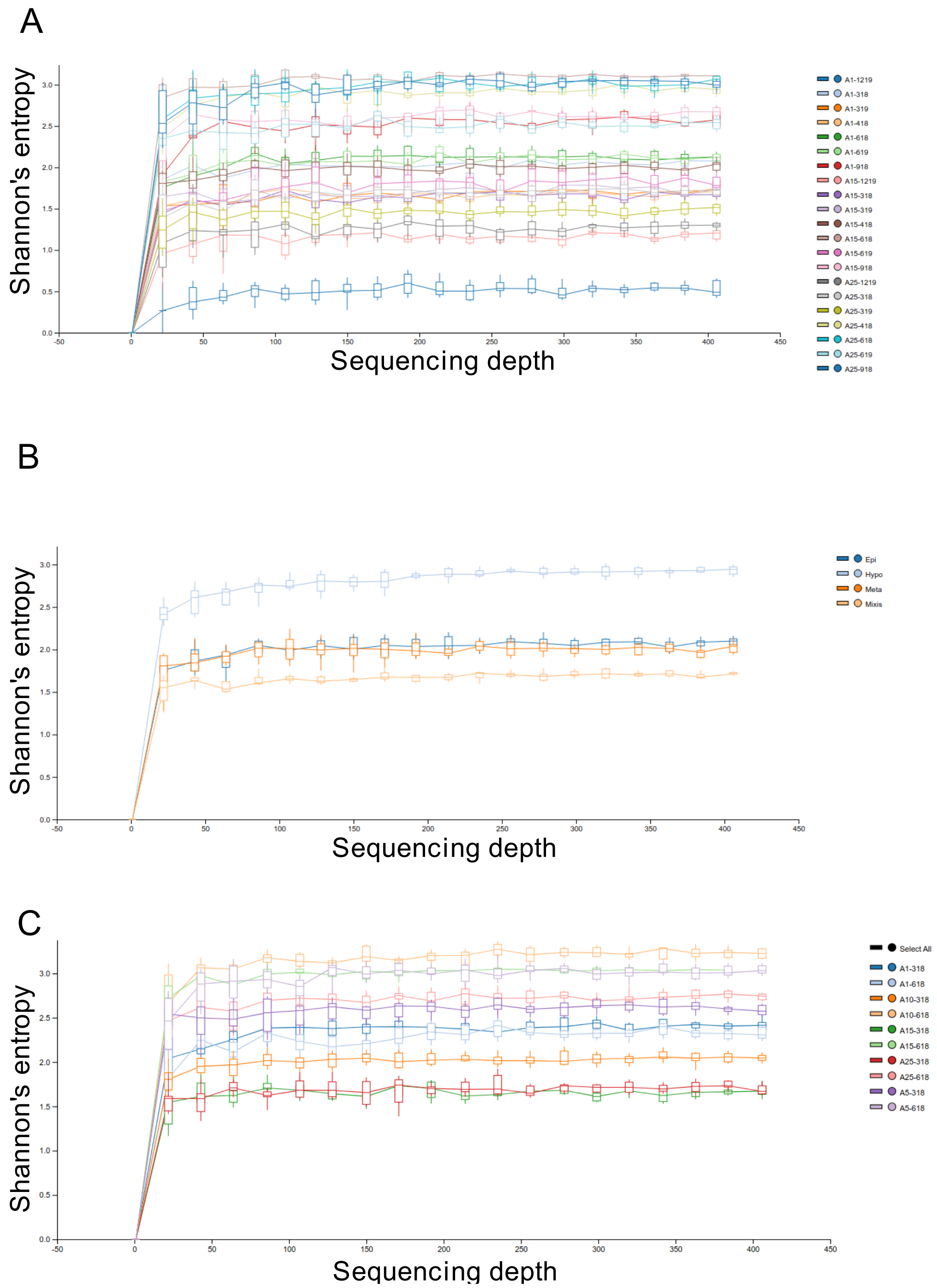


**Figure S4. Rarefaction curves for 16S amplicon-based Next generation Sequencing (NGS).** NGS was performed using *Legionella* spp. specific 16S primers. Raw data were pre-filtered to retain *Legionella* Amplicon Sequence Variants (ASVs), present at a minimal frequency of 20 reads across all samples, in a minimum of 2 samples. Rarefaction curves generated using the QIIME2 alpha-rarefaction function of the q2-diversity plugin, are presented for **(A)** each month-depth combination, **(B)** sampling layers (Epilimnion = Epi, Metalimnion = Meta, Hypolimnion = Hypo and mixing period = Mixis), and **(C)** for the 03.2018 and 06.2018 depth profiles**.**

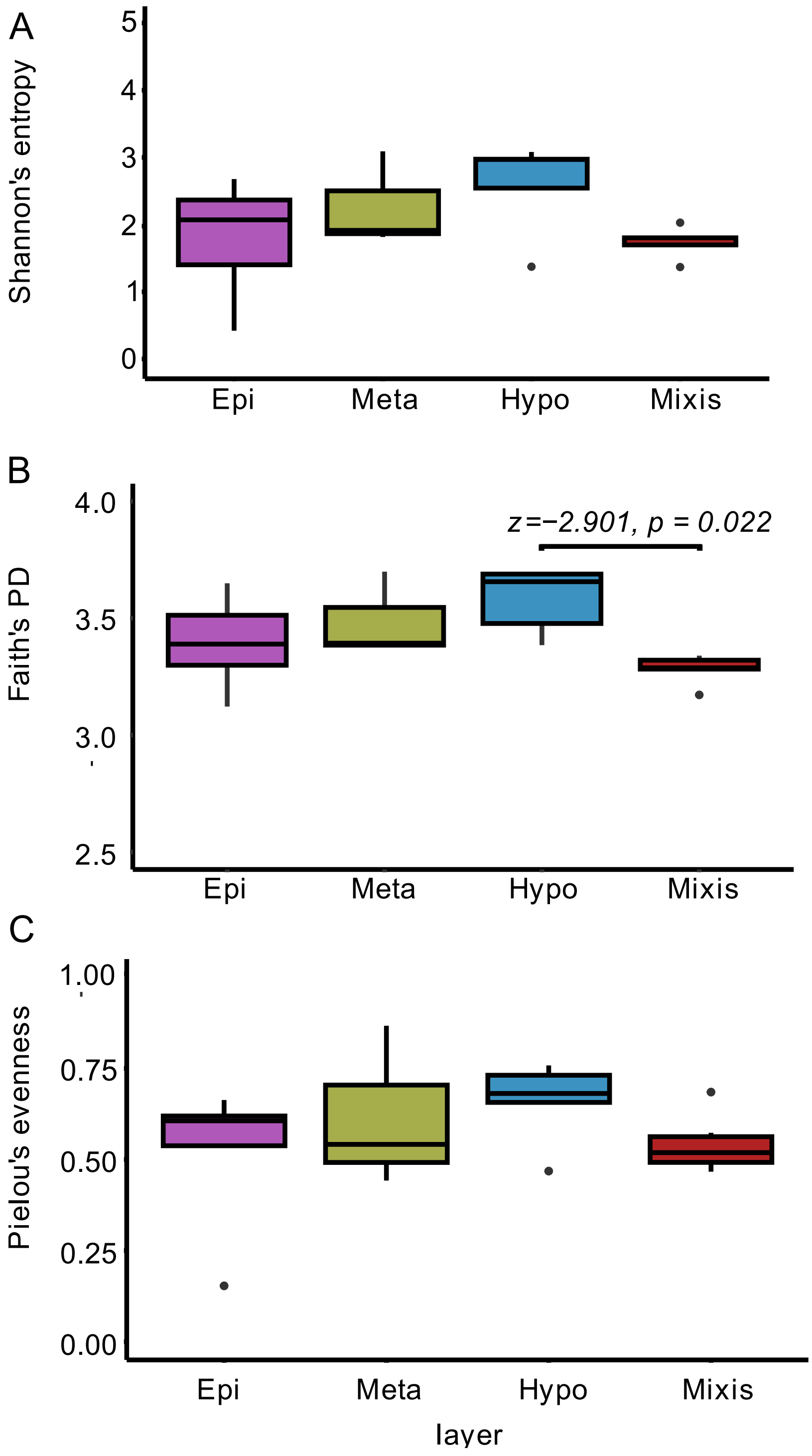


**Figure S5. Alpha diversity of *Legionella* spp. microbial population for the different sampled water column layers.** Next Generation Sequencing (NGS) was performed using *Legionella* spp. specific 16S primers. Raw data were pre-filtered to retain *Legionella* Amplicon Sequence Variants (ASVs), at a minimal frequency of 20 reads across all samples, in a minimum of 2 samples. **(A)** Shannon's entropy, **(B)** Faith's PD and **(C)** Pielou's evenness were included in the analysis. Kruskal-Wallis test results and post hoc via Wilcox tests are presented in Supplementary Tables S13 and S15.

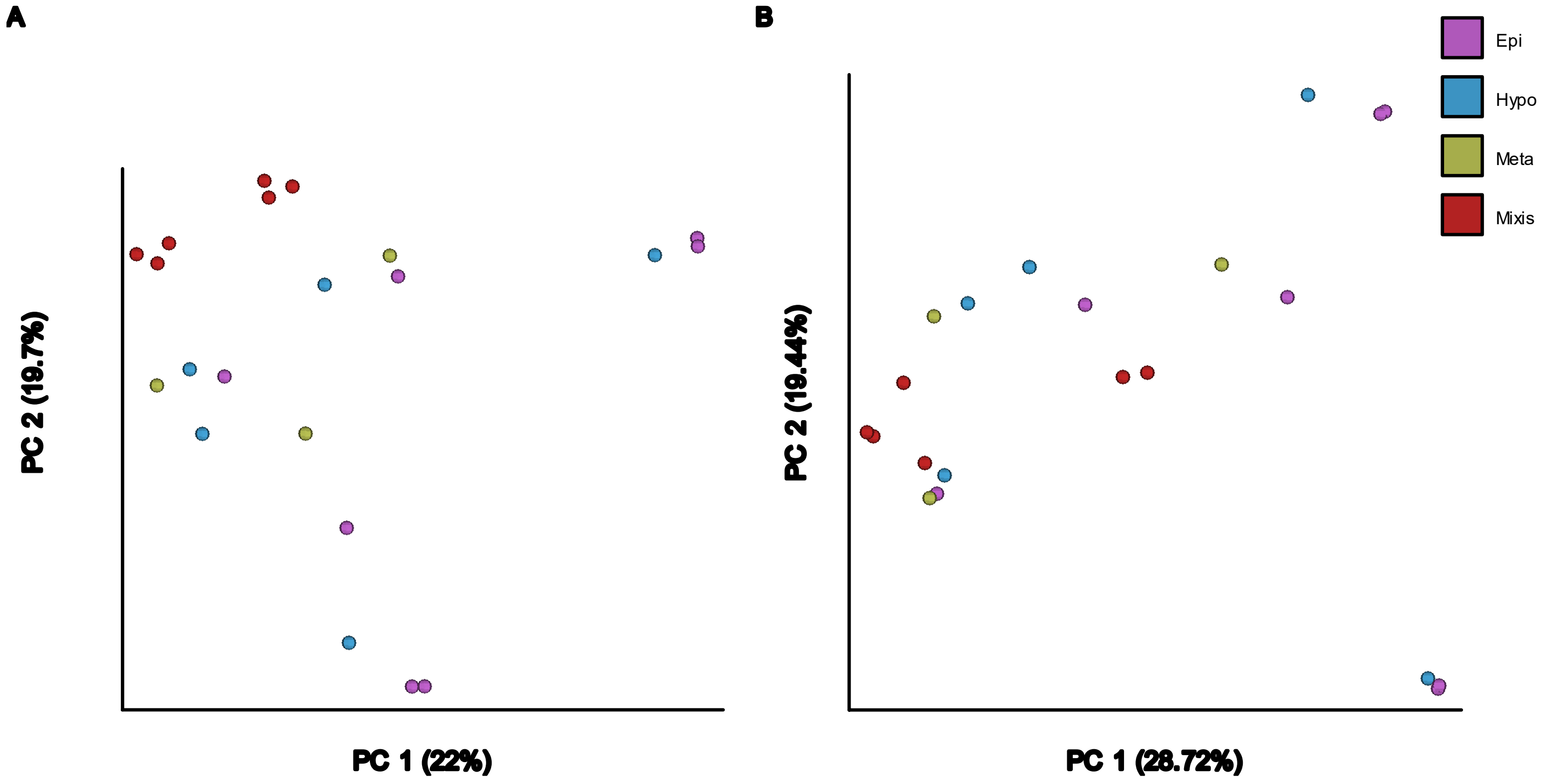


**Figure S6. Beta diversity of *Legionella* spp. microbial population for the different sampled water column layers.** Next Generation Sequencing (NGS) was performed using *Legionella* spp. specific 16S primers. Raw data were pre-filtered to retain *Legionella* Amplicon Sequence Variants (ASVs), at a minimal frequency of 20 reads across all samples, in a minimum of 2 samples. Beta diversity was assessed via **(A)** Jaccard and **(B)** Bray–Curtis. Statistical significance was tested via PERMANOVA and PERMDISP analyses (Tables S16-S17).

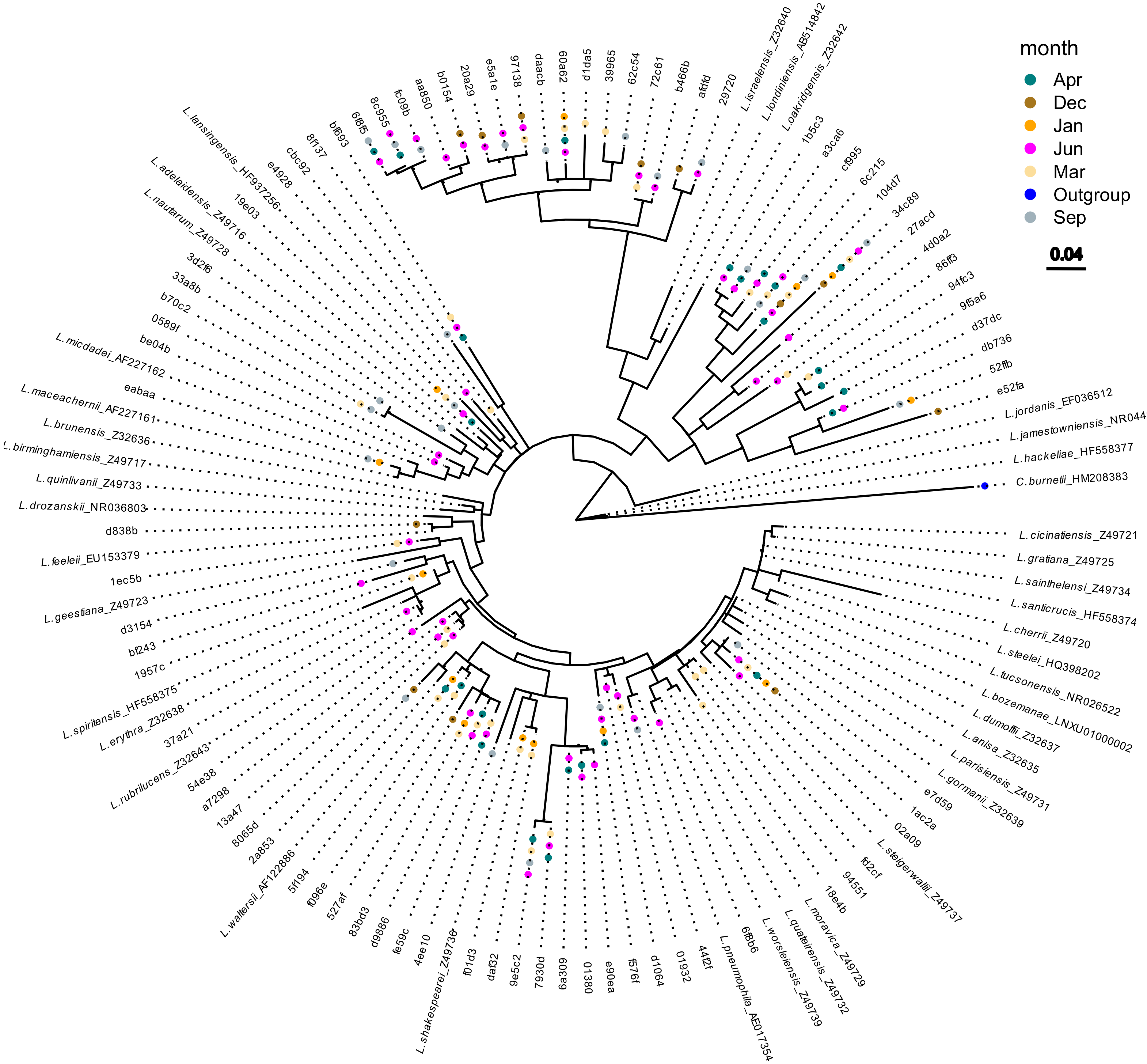


**Figure S7. Phylogenetic analysis of *Legionella* amplicon sequence variants (ASVs) according to month.** Phylogenetic tree was constructed alongside reference bacteria with corresponding NCBI accession-numbers (Supplementary Data 4). ASVs were filtered for minimum of 20 reads, in at least 2 samples. The evolutionary history of all trees was inferred using the Maximum-Likelihood method and General Time Reversible model (GTM), for the water spring clusters, and Hasegawa-Kishino-Yano (HKY) model. *Coxiella burnetii* was included as an outgroup. Tree with the highest log likelihood is presented

(-4775.69). A discrete Gamma distribution was used to model evolutionary rate differences among sites (5 categories (+G, parameter = 0.3503)). The rate variation model allowed for some sites to be evolutionarily invariable ([+I], 46.55% sites). The tree is drawn to scale, with branch lengths measured in the number of substitutions per site. Branches with bootstrap values above 50% are demoted by a black dot. The analysis involved 75 nucleotide sequences. There was a total of 380 positions in the final dataset. Evolutionary analyses were conducted in MEGA11.

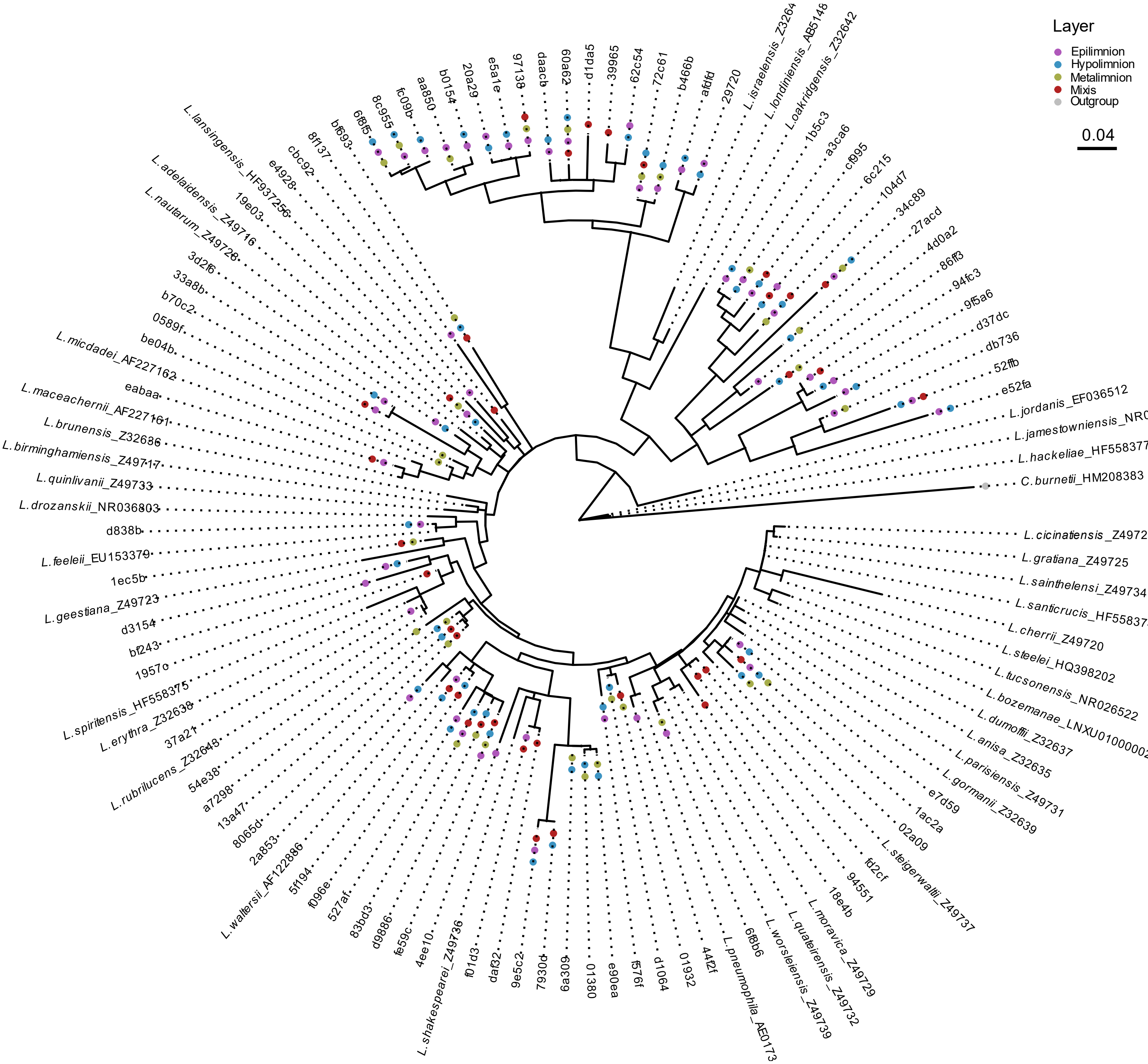


**Figure S8. Phylogenetic analysis of *Legionella* amplicon sequence variants (ASVs) according to sampling layer.** Phylogenetic tree was constructed alongside reference bacteria with corresponding NCBI accession-numbers (Supplementary Data 4). ASVs were filtered for minimum of 20 reads, in at least 2 samples. The evolutionary history of all trees was inferred using the Maximum-Likelihood method and General Time Reversible model (GTM), for the water spring clusters, and Hasegawa-Kishino-Yano (HKY) model. *Coxiella burnetii* was included as an outgroup. Tree with the highest log likelihood is presented (-4775.69). A discrete Gamma distribution was used to model evolutionary rate differences among sites (5 categories (+G, parameter = 0.3503)). The rate variation model allowed for some sites to be evolutionarily invariable ([+I], 46.55% sites). The tree is drawn to scale, with branch lengths measured in the number of substitutions per site. Branches with bootstrap values above 50% are demoted by a black dot. The analysis involved 75 nucleotide sequences. There was a total of 380 positions in the final dataset. Evolutionary analyses were conducted in MEGA11.

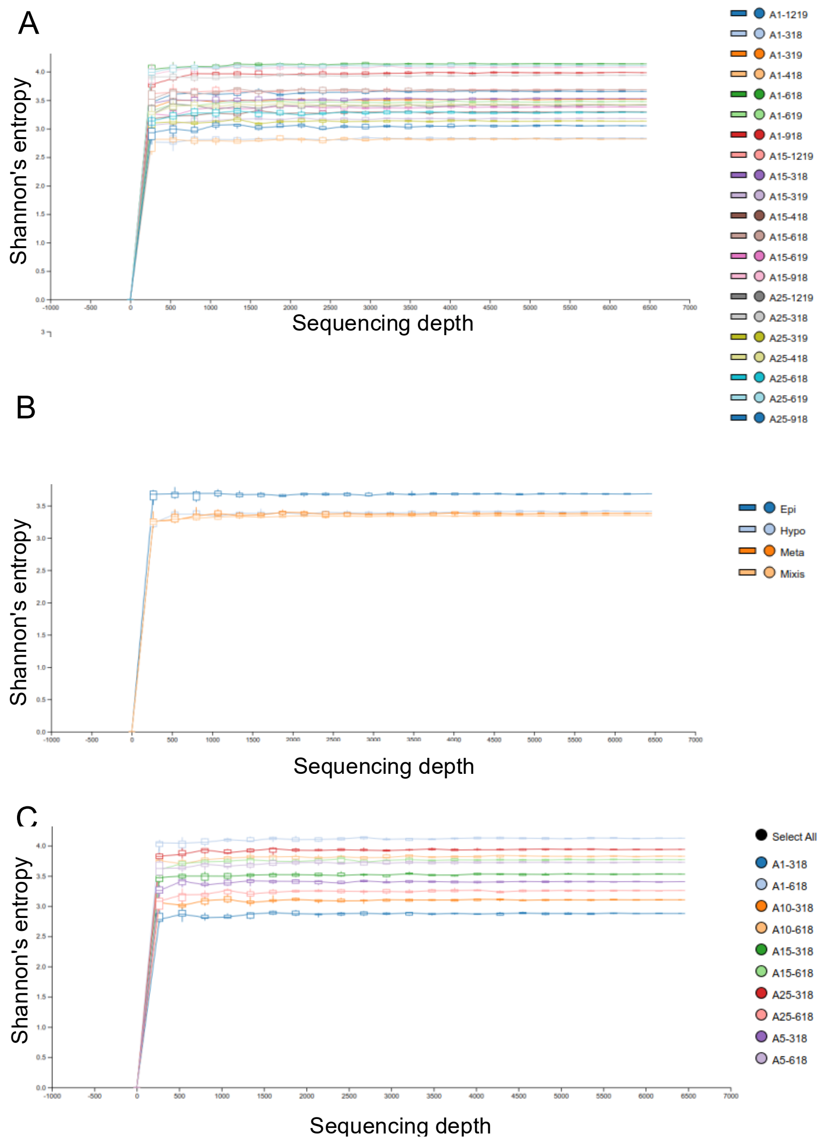


**Figure S9. Rarefaction curves for 18S amplicon-based Next generation Sequencing (NGS).** Next Generation Sequencing (NGS, n=28) was performed using universal protists primers, designed for the V9 region of the 18S rRNA gene. Raw data were pre-filtered to retain Amoebozoa, Cliophora (Ciliates), Percolozoa (Excavata) and Dinophyta (Dinoflagellates), at a minimal frequency of 20 reads across all samples, in a minimum of 2 samples. We included in the analyses water column samples from stratified layers: Epilimnion (Epi=purple, n=9), metalimnion (meta=pale-blue, n=4) hypolimnion (Hypo=mustered-green, n=5), and samples from the complete water column mixing period (mixis=maroon, n=10). Rarefaction curves generated using the QIIME2 alpha-rarefaction function of the q2-diversity plugin, are presented for **(A)** each month-depth combination, **(B)** sampling layers (Epilimnion = Epi, Metalimnion = Meta, Hypolimnion = Hypo and mixing period = Mixis), and **(C)** for the 03.2018 and 06.2018 depth profiles**.**

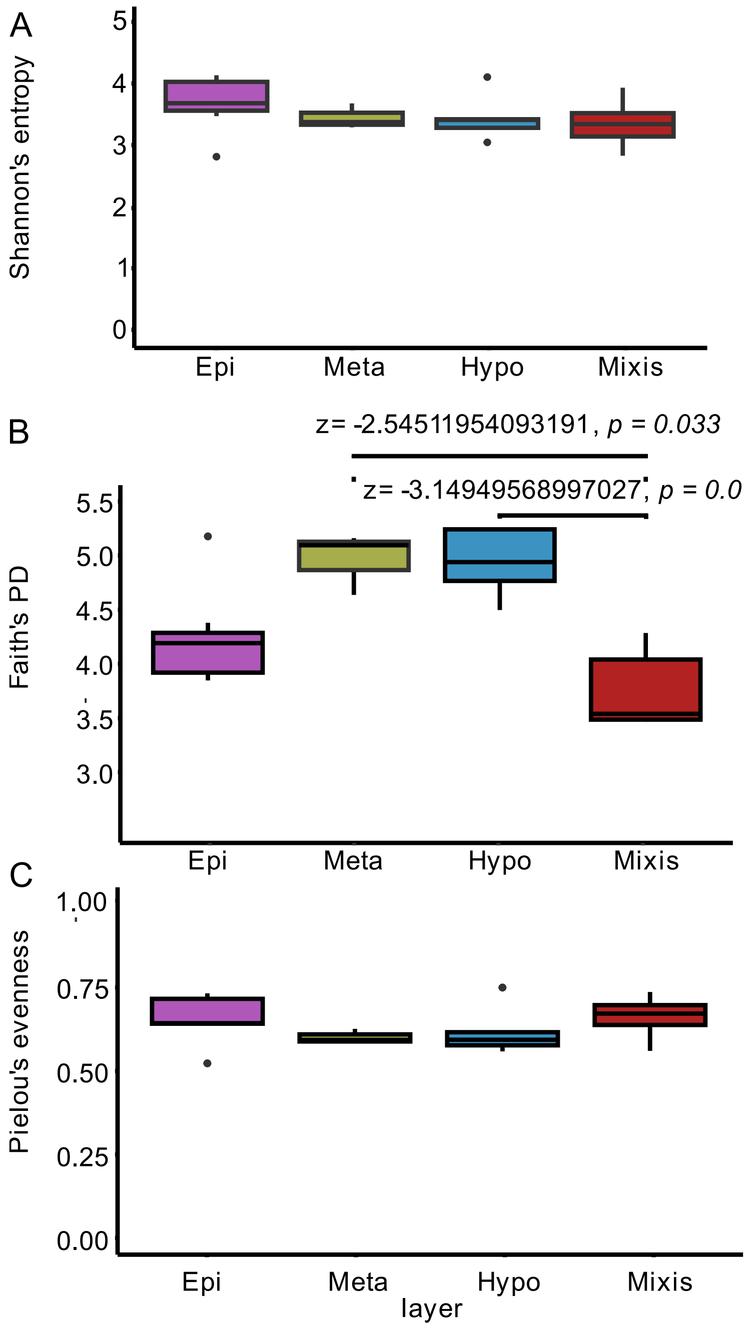


**Figure S10. Alpha diversity of *Legionella* spp. potential host's microbial population for the different sampled water column layers** Next Generation Sequencing (NGS, n=28) was performed using universal protists primers, designed for the V9 region of the 18S rRNA gene. Raw data were pre-filtered to retain Amoebozoa, Cliophora (Ciliates), Percolozoa (Excavata) and Dinophyta (Dinoflagellates), at a minimal frequency of 20 reads across all samples, in a minimum of 2 samples. **(A)** Shannon's entropy, **(B)** Faith's PD and **(C)** Pielou's evenness were included in the analysis. Kruskal-Wallis test results and post hoc via Wilcox tests are presented in Supplementary Tables S19-S20.

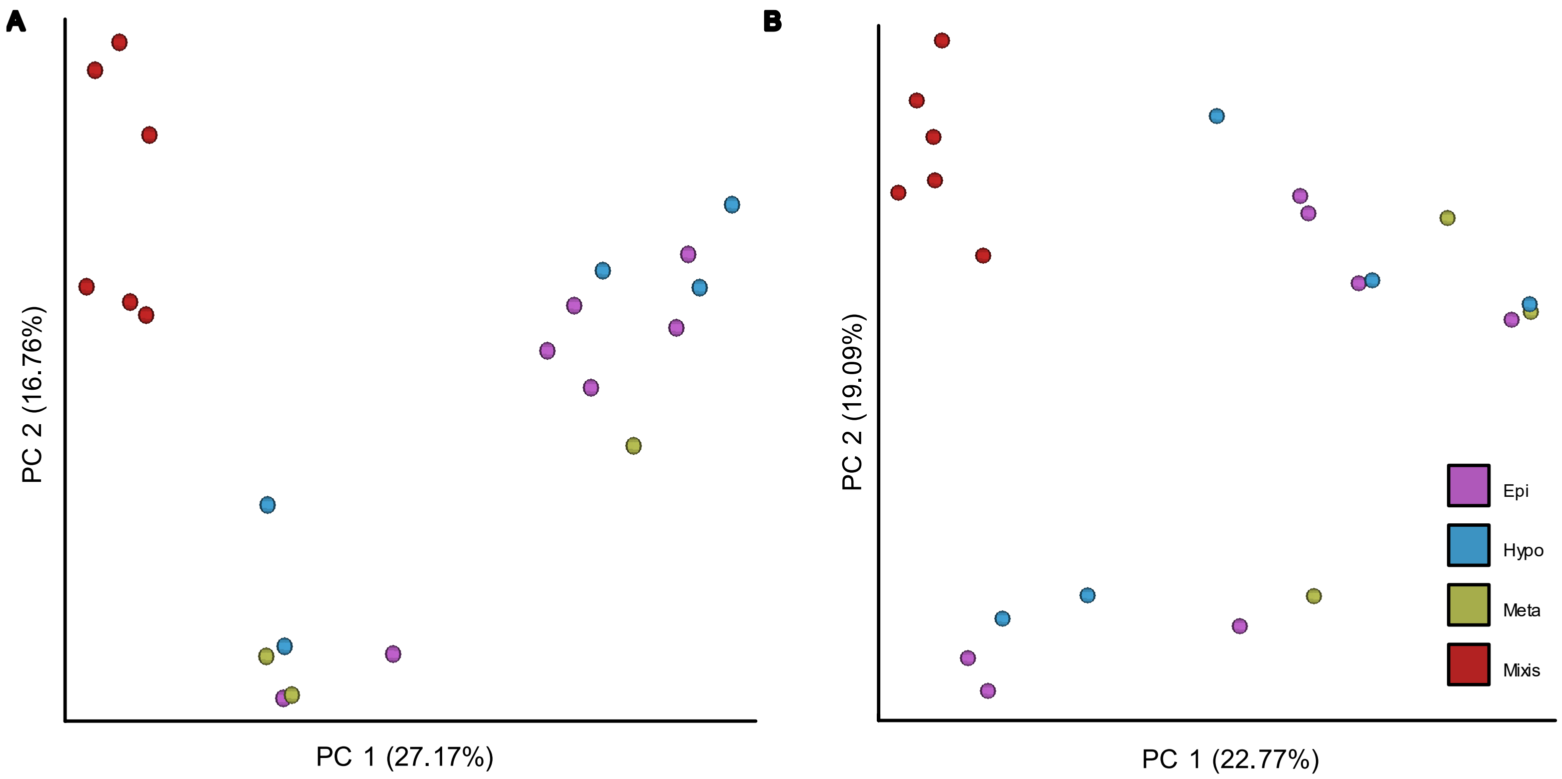


**Figure S11. Beta diversity of *Legionella* spp. potential host's microbial community composition for the different sampled water column layers.** Next Generation Sequencing (NGS, n=28) was performed using universal protists primers, designed for the V9 region of the 18S rRNA gene. Raw data were pre-filtered to retain Amoebozoa, Cliophora (Ciliates), Percolozoa (Excavata) and Dinophyta (Dinoflagellates), at a minimal frequency of 20 reads across all samples, in a minimum of 2 samples. Beta diversity was assessed via **(A)** Jaccard and **(B)** Bray–Curtis. Statistical significance was tested via PERMANOVA and PERMDISP analyses (Tables S21-S22).

**Table S1:** *Legionella* absolute abundance ANOVA Table (type II tests) for layer and post-hoc tukey test. Tested variable was layer, defined as Epilimnion (Epi), Metalimnion (Meta) and Hypolimnion (Hypo). Levene's test performed on the layer variable was not significant (df=2,60, F=0.883, *p*=0.418). p-value considered significant if < 0.05, marked bold with underline.

| ANOVA test | | | | Post hoc tukey test | | | | | |
|---|---|---|---|---|---|---|---|---|---|
| **F statistic** | **DFn** | **DFd** | **p-value** | **group1** | **group2** | **estimate** | **conf.low** | **conf.high** | **p.adj** |
| 3.67 | 2 | 60 | 0.031 | Epi | Hypo | 0.930 | 0.105 | 1.756 | **0.0235** |
| | | | | Epi | Meta | 0.330 | -0.839 | 1.498 | 0.777 |
| | | | | Hypo | Meta | -0.601 | -1.836 | 0.635 | 0.477 |

**Table S2:** Linear regression for depth profile March 2018, between Legionella count [cells / ml] to biotic and abiotic parameters in station A. p-value considered significant if < 0.05, marked bold with underline.

| Parameter (x) | Test statistic (S) | rho | p-value |
|---|---|---|---|
| **Temperature** | 68 | -0.943 | **0.005** |
| **DO** | 68 | -0.943 | **0.005** |
| **Chlorophyll** | 68 | -0.943 | **0.005** |
| **pH** | 68 | -0.943 | **0.005** |
| **Cl** | 6 | 0.841 | **0.036** |
| **Ptot** | 37 | -0.058 | 0.913 |
| **SO4** | 13 | 0.621 | 0.188 |
| **NH4** | 22 | 0.377 | 0.461 |

**Table S3:** Linear regression for depth profile June 2018, between Legionella count [cells / ml] to biotic and abiotic parameters in station A. p-value considered significant if < 0.05, marked bold with underline

| Parameter (x) | Test statistic (S) | rho | p-value |
|---|---|---|---|
| **Temperature** | 886.58 | -0.58 | **<u>0.022</u>** |
| **DO** | 847.71 | -0.51 | **<u>0.050</u>** |
| **Chlorophyll** | 640 | -0.14 | 0.612 |
| **pH** | 841.04 | -0.50 | 0.057 |
| **Cl** | 75.21 | 0.54 | 0.104 |
| **Ptot** | 77.85 | 0.07 | 0.863 |
| **SO4** | 70.96 | -0.27 | 0.562 |
| **NH4** | 74.90 | 0.55 | 0.102 |

**Table S4:** *Legionella* absolute abundance t-test table. Tested variable was stratification period, defined as Mixis (no stratification) and Stratified (into Epilimnion, Metalimnion and Hypolimnion layers).

| .y. | group1 | group2 | n1 | n2 | statistic | df | p-value |
|---|---|---|---|---|---|---|---|
| Leg_log | Mixis | Stratified | 21 | 63 | 0.831 | 40.339 | 0.411 |

Linear mixed model fit by maximum likelihood. t-tests use Satterthwaite's method ['lmerModLmerTest']

Formula: Logscale ~ sampling month + (1 | layer)

**Table S5:** Linear mixed model (LMM) Scaled residuals.

| Min | 1Q | Median | 3Q | Max |
|---|---|---|---|---|
| -2.244 | -0.426 | 0.000 | 0.426 | 2.245 |

**Table S6:** Linear mixed model (LMM) Type III Analysis of Variance Table with Satterthwaite's method.

| | Sum Sq | Mean Sq | NumDF | DenDF | F value | Pr(>F) |
|---|---|---|---|---|---|---|
| Sampling | 43.249 | 1.4914 | 29 | 34.134 | 11.934 | **7.67e-11** |

**Table S7:** Linear mixed model (LMM) Random effects.

| Groups | Name | Variance | Std.Dev. |
|---|---|---|---|
| layer | (Intercept) | 0.0619 | 0.249 |
| Residual | | 0.125 | 0.354 |

Number of obs: 54, groups: layer, 4

**Table S8**: Linear mixed model (LMM) fixed effects.

| | Estimate | Std. Error | df | t value | Pr(>\|t\|) |
|---|---|---|---|---|---|
| (Intercept) | 0.394 | 0.258 | 51.544 | 1.525 | 0.133 |
| Sep-19 | -2.14725 | 0.358664 | 52.83067 | -5.987 | **1.92E-07** |
| Mar-19 | 0.791035 | 0.373758 | 40.73305 | 2.116 | **0.040** |
| Aug-19 | -1.38576 | 0.358664 | 52.83067 | -3.864 | **0.000** |
| Jun-17 | 1.183747 | 0.436241 | 50.72908 | 2.714 | **0.009** |
| Mar-18 | 0.694493 | 0.373758 | 40.73305 | 1.858 | 0.070 |
| Dec-17 | 0.146691 | 0.357685 | 51.77189 | 0.41 | 0.683 |
| Sep-18 | -0.07032 | 0.35351 | 48.73002 | -0.199 | 0.843 |
| Jul-17 | 1.060382 | 0.436241 | 50.72908 | 2.431 | **0.019** |
| Aug-18 | -0.06603 | 0.35351 | 48.73002 | -0.187 | 0.853 |
| Feb-18 | -0.11711 | 0.373758 | 40.73305 | -0.313 | 0.756 |
| Jun-18 | -1.20111 | 0.358664 | 52.83067 | -3.349 | **0.002** |
| Nov-19 | -0.92257 | 0.35351 | 48.73002 | -2.61 | **0.012** |
| Jun-19 | 0.683054 | 0.358664 | 52.83067 | 1.904 | 0.062 |
| Oct-18 | -1.01827 | 0.35351 | 48.73002 | -2.88 | **0.006** |
| Jan-18 | -0.55449 | 0.373758 | 40.73305 | -1.484 | 0.146 |
| Oct-17 | 0.327579 | 0.436241 | 50.72908 | 0.751 | 0.456 |
| Oct-19 | -1.44656 | 0.35351 | 48.73002 | -4.092 | **0.000** |
| Dec-18 | -1.39213 | 0.357521 | 51.56521 | -3.894 | **0.000** |
| Jul-18 | -1.64884 | 0.35351 | 48.73002 | -4.664 | **0.000** |
| Feb-19 | -0.0301 | 0.373758 | 40.73305 | -0.081 | 0.936 |
| Sep-17 | -0.00908 | 0.436241 | 50.72908 | -0.021 | 0.983 |
| Nov-17 | -0.11408 | 0.436241 | 50.72908 | -0.261 | 0.795 |
| Jan-19 | -1.48763 | 0.373758 | 40.73305 | -3.98 | **0.000** |
| May-17 | 1.23806 | 0.436241 | 50.72908 | 2.838 | **0.007** |
| Apr-18 | 0.859024 | 0.358664 | 52.83067 | 2.395 | **0.020** |
| May-19 | -0.36069 | 0.358664 | 52.83067 | -1.006 | 0.319 |
| Nov-18 | -1.81907 | 0.35351 | 48.73002 | -5.146 | **0.000** |
| Apr-19 | -0.27443 | 0.373758 | 40.73305 | -0.734 | 0.467 |
| Jul-18 | -0.42923 | 0.358664 | 52.83067 | -1.197 | 0.237 |

**Table S9:** Correlations between the physicochemical parameters comprising the initial PCA. Correlations ≥ 0.7 p-value considered significant if < 0.05, marked bold with underline.

| Variable 1 | Variable 2 | cor | p-value |
|---|---|---|---|
| Temperature | Conductivity | -0.166 | 0.132 |
| Temperature | DO | 0.331 | 0.002 |
| Conductivity | DO | -0.106 | 0.337 |
| Temperature | Cl | 0.068 | 0.539 |
| **Conductivity** | **Cl** | **0.863** | **0.000** |
| DO | Cl | 0.041 | 0.710 |
| Temperature | $SO_4$ | 0.067 | 0.548 |
| Conductivity | $SO_4$ | 0.528 | 0.000 |
| DO | $SO_4$ | 0.385 | 0.000 |
| Cl | $SO_4$ | 0.597 | 0.000 |
| Temperature | Turbidity | 0.070 | 0.526 |
| Conductivity | Turbidity | -0.111 | 0.314 |
| DO | Turbidity | -0.088 | 0.424 |
| Cl | Turbidity | -0.102 | 0.358 |
| $SO_4$ | Turbidity | -0.167 | 0.131 |
| Temperature | pH | 0.638 | 0.000 |
| Conductivity | pH | -0.148 | 0.180 |
| DO | pH | 0.670 | 0.000 |
| Cl | pH | 0.010 | 0.925 |
| $SO_4$ | pH | 0.239 | 0.030 |
| Turbidity | pH | 0.194 | 0.077 |
| **Temperature** | **Alkalinity** | **-0.798** | **0.000** |
| Conductivity | Alkalinity | 0.115 | 0.297 |
| DO | Alkalinity | -0.558 | 0.000 |
| Cl | Alkalinity | -0.156 | 0.156 |
| $SO_4$ | Alkalinity | -0.333 | 0.002 |
| Turbidity | Alkalinity | 0.134 | 0.224 |
| pH | Alkalinity | -0.686 | 0.000 |
| **Temperature** | **Ca** | **-0.726** | **0.000** |

| | | | |
|---|---|---|---|
| Conductivity | Ca | 0.060 | 0.590 |
| DO | Ca | -0.659 | 0.000 |
| Cl | Ca | -0.205 | 0.062 |
| So4 | Ca | -0.363 | 0.001 |
| Turbidity | Ca | 0.012 | 0.915 |
| **pH** | **Ca** | **-0.766** | **0.000** |
| **Alkalinity** | **Ca** | **0.853** | **0.000** |
| Temperature | NH4 | -0.430 | 0.000 |
| Conductivity | NH4 | 0.003 | 0.975 |
| DO | NH4 | -0.466 | 0.000 |
| Cl | NH4 | -0.063 | 0.572 |
| $SO_4$ | NH4 | -0.393 | 0.000 |
| Turbidity | NH4 | 0.169 | 0.124 |
| pH | NH4 | -0.465 | 0.000 |
| Alkalinity | NH4 | 0.608 | 0.000 |
| Ca | NH4 | 0.588 | 0.000 |
| Temperature | Nitrit | -0.493 | 0.000 |
| Conductivity | Nitrit | -0.028 | 0.797 |
| DO | Nitrit | 0.284 | 0.009 |
| Cl | Nitrit | -0.140 | 0.205 |
| So4 | Nitrit | 0.216 | 0.050 |
| Turbidity | Nitrit | -0.204 | 0.063 |
| pH | Nitrit | -0.173 | 0.115 |
| Alkalinity | Nitrit | 0.163 | 0.138 |
| Ca | Nitrit | 0.151 | 0.171 |
| NH4 | Nitrit | -0.118 | 0.286 |
| Temperature | Ntot | -0.737 | 0.000 |
| Conductivity | Ntot | -0.081 | 0.462 |
| DO | Ntot | -0.204 | 0.063 |
| Cl | Ntot | -0.284 | 0.009 |
| So4 | Ntot | -0.215 | 0.051 |
| Turbidity | Ntot | 0.177 | 0.106 |

| | | | |
|---|---|---|---|
| pH | Ntot | -0.459 | 0.000 |
| **Alkalinity** | **Ntot** | **0.748** | **0.000** |
| Ca | Ntot | 0.537 | 0.000 |
| NH4 | Ntot | 0.379 | 0.000 |
| Nitrit | Ntot | 0.448 | 0.000 |
| Temperature | Ptot | -0.401 | 0.000 |
| Conductivity | Ptot | 0.067 | 0.544 |
| DO | Ptot | 0.015 | 0.889 |
| Cl | Ptot | -0.054 | 0.626 |
| So4 | Ptot | -0.097 | 0.383 |
| Turbidity | Ptot | 0.234 | 0.032 |
| pH | Ptot | -0.170 | 0.121 |
| Alkalinity | Ptot | 0.303 | 0.005 |
| Ca | Ptot | 0.192 | 0.080 |
| NH4 | Ptot | 0.179 | 0.103 |
| Nitrit | Ptot | 0.431 | 0.000 |
| Ntot | Ptot | 0.543 | 0.000 |
| Temperature | Chl | 0.618 | 0.000 |
| Conductivity | Chl | -0.181 | 0.107 |
| DO | Chl | 0.615 | 0.000 |
| Cl | Chl | 0.011 | 0.920 |
| So4 | Chl | 0.315 | 0.005 |
| Turbidity | Chl | 0.151 | 0.180 |
| pH | Chl | 0.673 | 0.000 |
| Alkalinity | Chl | -0.571 | 0.000 |
| Ca | Chl | -0.593 | 0.000 |
| NH4 | Chl | -0.349 | 0.001 |
| Nitrit | Chl | -0.084 | 0.458 |
| Ntot | Chl | -0.380 | 0.001 |
| Ptot | Chl | -0.114 | 0.313 |

**Table S10:** Loading scores of environmental variables included in the initial PCA analysis.

| Parameter | PC1 | PC2 |
|---|---|---|
| **Temperature** | -0.342 | 0.430 |
| **DO** | -0.439 | -0.170 |
| **Cl** | -0.037 | -0.216 |
| **So4** | -0.380 | -0.326 |
| **Turbidity** | 0.204 | 0.326 |
| **pH** | -0.371 | 0.270 |
| **NH4** | 0.438 | 0.174 |
| **Nitrit** | -0.005 | -0.577 |
| **Ptot** | 0.139 | -0.242 |
| **Chl** | -0.392 | 0.181 |

**Table S11:** Loading scores of environmental variables included in the final PCA analysis.

| Parameter | PC1 | PC2 |
|---|---|---|
| **Temperature** | -0.348 | 0.522 |
| **DO** | -0.453 | -0.167 |
| **$SO_4$** | -0.377 | -0.579 |
| **pH** | -0.407 | 0.408 |
| **$NH_4$** | 0.435 | 0.366 |
| **Chl** | -0.422 | 0.252 |

**Table S12**: Linear regression between *Legionella* count [cells / ml] to biotic and abiotic parameters in station A. Data presented for depths 1m, 15m and 25m - between May 2017 and December 2019 p-value considered significant if < 0.05, marked bold with underline.

| Parameter (x) | Test statistic (S) | rho | p-value |
|---|---|---|---|
| **Temperature** | 139279 | -0.41 | >0.001 |
| **Chlorophyll** | 121656 | -0.423 | >0.001 |
| **Ptot** | 110478 | -0.12 | 0.283 |
| **Nitrate** | 35986 | 0.176 | 0.163 |
| **NH4** | 95231 | 0.035 | 0.746 |
| **pH** | 132548 | -0.34 | 0.0014 |
| **DO** | 115237 | -0.17 | 0.13 |
| **Dinoflagellates** | 23043 | 0.12 | 0.381 |

**Table S13:** Kruskal-Wallis tests of 16S NGS alpha diversity between the different months, and layers.

| Comparison | Alpha index | H statistic | df | p-value |
|---|---|---|---|---|
| **Month** | **Shannon entropy** | 13.96 | 4 | **0.007** |
| | **Faith's PD** | 11.87 | 4 | **0.018** |
| | **Pielou's evenness** | 8.26 | 4 | 0.082 |
| **Layer** | **Shannon entropy** | 5.2 | 3 | 0.16 |
| | **Faith PD** | 9.64 | 3 | **0.021** |
| | **Pielou evenness** | 3.22 | 3 | 0.36 |

**Table S14:** Post hoc via Dunn's tests of 16S NGS alpha diversity between the different months (March, April, June, September, December).

| Alpha_index | group1 | group2 | n1 | n2 | statistic | p-value | p.adj |
|---|---|---|---|---|---|---|---|
| Faith pd | Apr | Dec | 3 | 3 | -1.053 | 0.292 | 0.418 |
| | Apr | Jun | 3 | 6 | 0.684 | 0.494 | 0.618 |
| | Apr | Mar | 3 | 6 | -1.481 | 0.138 | 0.277 |
| | Apr | Sep | 3 | 3 | 1.053 | 0.292 | 0.418 |
| | Dec | Jun | 3 | 6 | 1.899 | 0.058 | 0.144 |
| | Dec | Mar | 3 | 6 | -0.266 | 0.79 | 0.79 |
| | Dec | Sep | 3 | 3 | 2.105 | 0.035 | 0.118 |
| | Jun | Mar | 6 | 6 | -2.652 | 0.008 | **0.04** |
| | Jun | Sep | 6 | 3 | 0.532 | 0.595 | 0.661 |
| | Mar | Sep | 6 | 3 | 2.697 | 0.007 | **0.04** |
| Shannon entropy | Apr | Dec | 3 | 3 | -1.842 | 0.065 | 0.131 |
| | Apr | Jun | 3 | 6 | 0.798 | 0.425 | 0.472 |
| | Apr | Mar | 3 | 6 | -0.95 | 0.342 | 0.428 |
| | Apr | Sep | 3 | 3 | 1.184 | 0.236 | 0.341 |
| | Dec | Jun | 3 | 6 | 2.925 | 0.003 | **0.017** |
| | Dec | Mar | 3 | 6 | 1.178 | 0.239 | 0.341 |
| | Dec | Sep | 3 | 3 | 3.027 | 0.002 | **0.017** |
| | Jun | Mar | 6 | 6 | -2.14 | 0.032 | 0.081 |
| | Jun | Sep | 6 | 3 | 0.57 | 0.569 | 0.569 |
| | Mar | Sep | 6 | 3 | 2.317 | 0.02 | 0.068 |

**Table S15:** Post hoc via Dunn's tests of 16S NGS alpha diversity between the different layers (Epilimnion Metalimnion, Hypolimnion, mixis).

| Alpha index | group1 | group2 | n1 | n2 | statistic | p-value | p.adj |
|---|---|---|---|---|---|---|---|
| **Faith pd** | Epi | Hypo | 7 | 5 | 1.68 | 0.09 | 0.18 |
| | Epi | Meta | 7 | 3 | 1.02 | 0.31 | 0.37 |
| | Epi | Mixis | 7 | 6 | -1.39 | 0.17 | 0.25 |
| | Hypo | Meta | 5 | 3 | -0.38 | 0.70 | 0.70 |
| | Hypo | Mixis | 5 | 6 | -2.90 | 0.00 | 0.02 |
| | Meta | Mixis | 3 | 6 | -2.09 | 0.04 | 0.11 |

**Table S16:** PERMANOVA and PERMDISP results of 16S NGS beta diversity between the different months, and layers. Number of permutations = 999.

| Beta index | Test | Comparison | df | n | test statistic | p-value |
|---|---|---|---|---|---|---|
| **Jaccard** | **PERMANOVA** | Month | 5 | 21 | 5.18 | **0.001** |
| | | Layer | 4 | 21 | 1.78 | **0.014** |
| **Bray Curtis** | | Month | 5 | 21 | 5.14 | **0.001** |
| | | Layer | 4 | 21 | 1.5 | 0.086 |
| **Jaccard** | **PERMDISP** | Month | 5 | 21 | 7.63 | **0.005** |
| | | Layer | 4 | 21 | 3.87 | **0.018** |
| **Bray Curtis** | | Month | 5 | 21 | 2.95 | **0.05** |
| | | Layer | 4 | 21 | 1.68 | 0.116 |

**Table S17:** PERMANOVA and PERMDISP post hoc pairwise comparison results of 16S NGS beta diversity for month and layer. Number of permutations = 999.

| Bata index | Test | Comparison | Group 1 | Group 2 | n | pseudo-F | p-value | q-value |
|---|---|---|---|---|---|---|---|---|
| **Jaccard** | **PERMANOVA** | **Month** | Apr | Dec | 6 | 11.461 | 0.088 | 0.104 |
| | | | Apr | Jun | 9 | 1.912 | 0.054 | 0.077 |
| | | | Apr | Mar | 9 | 3.590 | 0.03 | **0.050** |
| | | | Apr | Sep | 6 | 8.205 | 0.104 | 0.104 |
| | | | Dec | Jun | 9 | 4.473 | 0.012 | **0.032** |
| | | | Dec | Mar | 9 | 9.837 | 0.013 | **0.032** |
| | | | Dec | Sep | 6 | 20.487 | 0.099 | 0.104 |
| | | | Jun | Mar | 12 | 3.416 | 0.006 | **0.032** |
| | | | Jun | Sep | 9 | 3.523 | 0.016 | **0.032** |
| | | | Mar | Sep | 9 | 9.322 | 0.009 | **0.032** |
| **Bray Curtis** | **PERMANOVA** | **Month** | Apr | Dec | 6 | 16.920 | 0.089 | 0.132 |
| | | | Apr | Jun | 9 | 1.854 | 0.107 | 0.132 |
| | | | Apr | Mar | 9 | 2.864 | 0.064 | 0.128 |
| | | | Apr | Sep | 6 | 10.623 | 0.119 | 0.132 |
| | | | Dec | Jun | 9 | 5.347 | 0.01 | **0.050** |
| | | | Dec | Mar | 9 | 9.988 | 0.015 | **0.050** |
| | | | Dec | Sep | 6 | 32.417 | 0.098 | 0.132 |
| | | | Jun | Mar | 12 | 1.434 | 0.236 | 0.236 |
| | | | Jun | Sep | 9 | 4.411 | 0.027 | 0.068 |
| | | | Mar | Sep | 9 | 8.373 | 0.008 | **0.050** |
| **Jaccard** | **PERMDISP** | **Month** | Mar | Dec | 9 | 5.606 | 0.032 | 0.055 |
| | | | Mar | Apr | 9 | 0.549 | 0.428 | 0.428 |
| | | | Mar | Jun | 12 | 6.078 | 0.002 | **0.020** |
| | | | Mar | Sep | 9 | 3.650 | 0.033 | 0.055 |
| | | | Apr | Dec | 6 | 4.272 | 0.1 | 0.125 |
| | | | Apr | Jun | 9 | 18.968 | 0.01 | **0.050** |
| | | | Apr | Sep | 6 | 1.634 | 0.062 | 0.089 |
| | | | Jun | Dec | 9 | 41.956 | 0.016 | **0.050** |
| | | | Jun | Sep | 9 | 22.282 | 0.02 | **0.050** |
| | | | Sep | Dec | 6 | 0.030 | 0.243 | 0.270 |
| | | | Mar | Dec | 9 | 4.874 | 0.013 | 0.057 |
| | | | Mar | Apr | 9 | 0.585 | 0.353 | 0.353 |

| | | | | | | | | |
|---|---|---|---|---|---|---|---|---|
| **Bray Curtis** | **PERMDISP** | **Month** | Mar | Jun | 12 | 0.916 | 0.322 | 0.353 |
| | | | Mar | Sep | 9 | 2.021 | 0.081 | 0.129 |
| | | | Apr | Dec | 6 | 1.499 | 0.044 | 0.104 |
| | | | Apr | Jun | 9 | 2.794 | 0.052 | 0.104 |
| | | | Apr | Sep | 6 | 0.249 | 0.117 | 0.146 |
| | | | Jun | Dec | 9 | 12.185 | 0.017 | 0.057 |
| | | | Jun | Sep | 9 | 5.962 | 0.01 | 0.057 |
| | | | Sep | Dec | 6 | 0.600 | 0.09 | 0.129 |
| **Jaccard** | **PERMANOVA** | **Layer** | Epi | Hypo | 12 | 0.818 | 0.543 | 0.652 |
| | | | Epi | Meta | 10 | 0.932 | 0.445 | 0.652 |
| | | | Epi | Mixis | 13 | 3.592 | 0.001 | **0.006** |
| | | | Hypo | Meta | 8 | 0.450 | 0.983 | 0.983 |
| | | | Hypo | Mixis | 11 | 3.054 | 0.004 | **0.012** |
| | | | Meta | Mixis | 9 | 2.163 | 0.059 | 0.118 |
| **Bray Curtis** | **PERMANOVA** | **Layer** | Epi | Hypo | 12 | 0.991 | 0.448 | 0.538 |
| | | | Epi | Meta | 10 | 1.123 | 0.343 | 0.515 |
| | | | Epi | Mixis | 13 | 3.054 | 0.011 | 0.066 |
| | | | Hypo | Meta | 8 | 0.570 | 0.812 | 0.812 |
| | | | Hypo | Mixis | 11 | 1.639 | 0.152 | 0.456 |
| | | | Meta | Mixis | 9 | 1.281 | 0.295 | 0.515 |
| **Jaccard** | **PERMDISP** | **Layer** | Epi | Hypo | 12 | 1.979 | 0.031 | 0.093 |
| | | | Epi | Meta | 10 | 3.358 | 0.236 | 0.283 |
| | | | Epi | Mixis | 13 | 14.019 | 0.001 | **0.006** |
| | | | Hypo | Meta | 8 | 0.007 | 0.94 | 0.940 |
| | | | Hypo | Mixis | 11 | 2.406 | 0.12 | 0.240 |
| | | | Meta | Mixis | 9 | 2.115 | 0.188 | 0.282 |
| **Bray Curtis** | **PERMDISP** | **Layer** | Epi | Hypo | 12 | 1.474 | 0.059 | 0.118 |
| | | | Epi | Meta | 10 | 5.815 | 0.043 | 0.118 |
| | | | Epi | Mixis | 13 | 5.575 | 0.007 | **0.042** |
| | | | Hypo | Meta | 8 | 0.037 | 0.78 | 0.780 |
| | | | Hypo | Mixis | 11 | 0.608 | 0.398 | 0.597 |
| | | | Meta | Mixis | 9 | 0.305 | 0.503 | 0.604 |

**Table S18:** ADONIS test results of 16S NGS beta diversity with selected environmental variables. Number of permutations = 999.

| Beta index | Variable | df | SumsOfSqs | MeanSqs | F.Model | R2 | Pr(>F) |
|---|---|---|---|---|---|---|---|
| Jaccard | **Temperature** | 1 | 0.859 | 0.859 | 7.217 | 0.116 | **0.001** |
| | **Cl** | 1 | 1.047 | 1.047 | 8.791 | 0.142 | **0.001** |
| | **DO** | 1 | 0.640 | 0.640 | 5.378 | 0.087 | **0.001** |
| | **pH** | 1 | 0.621 | 0.621 | 5.212 | 0.084 | **0.001** |
| | **Ptot** | 1 | 0.416 | 0.416 | 3.496 | 0.056 | **0.001** |
| | **NH4** | 1 | 0.465 | 0.465 | 3.907 | 0.063 | **0.001** |
| | **month** | 4 | 2.090 | 0.522 | 4.388 | 0.283 | **0.001** |
| | **layer** | 2 | 0.297 | 0.149 | 1.248 | 0.040 | 0.241 |
| | **Residuals** | 8 | 0.953 | 0.119 | NA | 0.129 | NA |
| | **Total** | 20 | 7.389 | NA | NA | 1.000 | NA |
| Bray Curtis | **Temperature** | 1 | 0.922 | 0.922 | 11.530 | 0.125 | **0.001** |
| | **Cl** | 1 | 1.020 | 1.020 | 12.759 | 0.138 | **0.001** |
| | **DO** | 1 | 0.325 | 0.325 | 4.070 | 0.044 | **0.002** |
| | **pH** | 1 | 0.823 | 0.823 | 10.294 | 0.112 | **0.001** |
| | **Ptot** | 1 | 0.271 | 0.271 | 3.386 | 0.037 | **0.003** |
| | **NH4** | 1 | 0.504 | 0.504 | 6.307 | 0.068 | **0.001** |
| | **month** | 4 | 2.384 | 0.596 | 7.456 | 0.323 | **0.001** |
| | **layer** | 2 | 0.487 | 0.243 | 3.044 | 0.066 | **0.002** |
| | **Residuals** | 8 | 0.639 | 0.080 | NA | 0.087 | NA |
| | **Total** | 20 | 7.375 | NA | NA | 1.000 | NA |

Variables were chosen based on *Legionella* spp. known factors influencing growth and on the principal component analysis (Fig. 2A and S2 AND Table S8).

**Table S19:** Kruskal-Wallis tests of 18S NGS alpha diversity between the different months, and layers.

| Comparison | Alpha index | H statistic | df | p-value |
|---|---|---|---|---|
| **Month** | **Shannon entropy** | 4.389 | 4 | 0.355 |
| | **Faith's PD** | 8.545 | 4 | 0.073 |
| | **Pielou's evenness** | 4.18 | 4 | 0.382 |
| **Layer** | **Shannon entropy** | 3.03 | 3 | 0.387 |
| | **Faith PD** | 12.64 | 3 | **0.005** |
| | **Pielou evenness** | 3.7 | 3 | 0.295 |

**Table S20:** Post hoc via Wilcox tests of 18S NGS alpha diversity (Faith pd) between the different layers (Epilimnion Metalimnion, Hypolimnion, mixis).

| Alpha_index | group1 | group2 | n1 | n2 | statistic | p-value | p.adj |
|---|---|---|---|---|---|---|---|
| **Faith pd** | Epi | Hypo | 7 | 5 | 2.084 | 0.037 | 0.074 |
| | Epi | Meta | 7 | 3 | 1.613 | 0.107 | 0.160 |
| | Epi | Mixis | 7 | 6 | -1.235 | 0.217 | 0.260 |
| | Hypo | Meta | 5 | 3 | -0.147 | 0.883 | 0.883 |
| | Hypo | Mixis | 5 | 6 | -3.149 | 0.002 | 0.010 |
| | Meta | Mixis | 3 | 6 | -2.545 | 0.011 | 0.033 |

**Table S21:** PERMANOVA and PERMDISP results of 18S NGS beta diversity between the different months, and layers. Number of permutations = 999.

| Beta index | Test | Comparison | df | n | test statistic | p-value |
|---|---|---|---|---|---|---|
| **Jaccard** | **PERMANOVA** | Month | 5 | 21 | 4.523 | **0.001** |
| **Jaccard** | **PERMANOVA** | Layer | 4 | 21 | 2.985 | **0.001** |
| **Bray Curtis** | **PERMANOVA** | Month | 5 | 21 | 3.450 | **0.001** |
| **Bray Curtis** | **PERMANOVA** | Layer | 4 | 21 | 2.435 | **0.001** |
| **Jaccard** | **PERMDISP** | Month | 5 | 21 | 2.99 | **0.019** |
| **Jaccard** | **PERMDISP** | Layer | 4 | 21 | 2.58 | **0.04** |
| **Bray Curtis** | **PERMDISP** | Month | 5 | 21 | 5.6 | **0.013** |
| **Bray Curtis** | **PERMDISP** | Layer | 4 | 21 | 2.12 | 0.092 |

**Table S22:** PERMANOVA and PERMDISP post hoc pairwise comparison results of 18S NGS beta diversity for month and layer. Number of permutations = 999.

| Bata index | Test | Comparison | Group 1 | Group 2 | n | pseudo-F | p-value | q-value |
|---|---|---|---|---|---|---|---|---|
| **Jaccard** | **PERMANOVA** | **Month** | Apr | Dec | 6 | 8.854 | 0.106 | 0.106 |
| | | | Apr | Jun | 9 | 2.214 | 0.021 | **0.042** |
| | | | Apr | Mar | 9 | 5.098 | 0.011 | **0.04** |
| | | | Apr | Sep | 6 | 5.965 | 0.099 | 0.106 |
| | | | Dec | Jun | 9 | 3.184 | 0.037 | 0.053 |
| | | | Dec | Mar | 9 | 7.393 | 0.02 | **0.042** |
| | | | Dec | Sep | 6 | 4.536 | 0.096 | 0.106 |
| | | | Jun | Mar | 12 | 4.772 | 0.005 | **0.04** |
| | | | Jun | Sep | 9 | 2.418 | 0.034 | 0.053 |
| | | | Mar | Sep | 9 | 6.579 | 0.012 | **0.04** |
| **Bray Curtis** | **PERMANOVA** | **Month** | Apr | Dec | 6 | 13.753 | 0.087 | 0.103 |
| | | | Apr | Jun | 9 | 1.548 | 0.082 | 0.103 |
| | | | Apr | Mar | 9 | 5.763 | 0.008 | **0.035** |
| | | | Apr | Sep | 6 | 6.435 | 0.093 | 0.103 |
| | | | Dec | Jun | 9 | 2.155 | 0.037 | 0.062 |
| | | | Dec | Mar | 9 | 4.328 | 0.014 | **0.035** |
| | | | Dec | Sep | 6 | 4.460 | 0.105 | 0.105 |
| | | | Jun | Mar | 12 | 3.232 | 0.002 | **0.020** |
| | | | Jun | Sep | 9 | 2.103 | 0.034 | 0.062 |
| | | | Mar | Sep | 9 | 3.422 | 0.012 | **0.035** |
| **Jaccard** | **PERMDISP** | **Month** | Apr | Dec | 6 | 0.335 | 0.103 | 0.147 |
| | | | Apr | Jun | 9 | 11.028 | 0.018 | **0.045** |
| | | | Apr | Mar | 9 | 4.413 | 0.025 | **0.050** |
| | | | Apr | Sep | 6 | 0.204 | 0.151 | 0.189 |
| | | | Dec | Jun | 9 | 15.065 | 0.012 | **0.045** |
| | | | Dec | Mar | 9 | 7.894 | 0.017 | **0.045** |
| | | | Dec | Sep | 6 | 0.659 | 0.05 | 0.083 |
| | | | Jun | Mar | 12 | 4.563 | 0.011 | **0.045** |
| | | | Jun | Sep | 9 | 1.598 | 0.199 | 0.221 |
| | | | Mar | Sep | 9 | 0.248 | 0.37 | 0.370 |
| | | | Apr | Dec | 6 | 0.002 | 0.820 | 0.820 |
| | | | Apr | Jun | 9 | 27.050 | 0.010 | **0.033** |

| | | | | | | | | |
|---|---|---|---|---|---|---|---|---|
| **Bray Curtis** | **PERMDISP** | **Month** | Apr | Mar | 9 | 23.967 | 0.010 | **0.033** |
| | | | Apr | Sep | 6 | 2.748 | 0.095 | 0.172 |
| | | | Dec | Jun | 9 | 16.647 | 0.010 | **0.033** |
| | | | Dec | Mar | 9 | 14.723 | 0.021 | 0.053 |
| | | | Dec | Sep | 6 | 2.269 | 0.103 | 0.172 |
| | | | Jun | Mar | 12 | 0.153 | 0.637 | 0.796 |
| | | | Jun | Sep | 9 | 0.346 | 0.539 | 0.770 |
| | | | Mar | Sep | 9 | 0.156 | 0.764 | 0.820 |
| **Jaccard** | **PERMANOVA** | **Layer** | Epi | Hypo | 12 | 1.889 | 0.059 | 0.0885 |
| | | | Epi | Meta | 10 | 1.345 | 0.192 | 0.2304 |
| | | | Epi | Mixis | 13 | 5.658 | 0.001 | **0.006** |
| | | | Hypo | Meta | 8 | 1.024 | 0.322 | 0.322 |
| | | | Hypo | Mixis | 11 | 4.268 | 0.002 | **0.006** |
| | | | Meta | Mixis | 9 | 3.677 | 0.017 | **0.034** |
| **Bray Curtis** | **PERMANOVA** | **Layer** | Epi | Hypo | 12 | 1.498 | 0.13 | 0.195 |
| | | | Epi | Meta | 10 | 1.292 | 0.265 | 0.318 |
| | | | Epi | Mixis | 13 | 3.751 | 0.003 | **0.009** |
| | | | Hypo | Meta | 8 | 1.086 | 0.329 | 0.329 |
| | | | Hypo | Mixis | 11 | 2.957 | 0.001 | **0.006** |
| | | | Meta | Mixis | 9 | 3.850 | 0.012 | **0.024** |
| **Jaccard** | **PERMDISP** | **Layer** | Epi | Hypo | 12 | 0.984 | 0.194 | 0.291 |
| | | | Epi | Meta | 10 | 1.148 | 0.451 | 0.541 |
| | | | Epi | Mixis | 13 | 5.795 | 0.001 | **0.006** |
| | | | Hypo | Meta | 8 | 1.845 | 0.109 | 0.218 |
| | | | Hypo | Mixis | 11 | 8.238 | 0.022 | 0.066 |
| | | | Meta | Mixis | 9 | 0.053 | 0.785 | 0.785 |
| **Bray Curtis** | **PERMDISP** | **Layer** | Epi | Hypo | 12 | 0.677 | 0.337 | 0.491 |
| | | | Epi | Meta | 10 | 3.853 | 0.149 | 0.298 |
| | | | Epi | Mixis | 13 | 0.011 | 0.889 | 0.889 |
| | | | Hypo | Meta | 8 | 3.677 | 0.113 | 0.298 |
| | | | Hypo | Mixis | 11 | 0.602 | 0.409 | 0.491 |
| | | | Meta | Mixis | 9 | 2.666 | 0.088 | 0.298 |
